# Enhanced Branched-Chain Amino Acid Metabolism Improves Age-Related Reproduction in *C. elegans*

**DOI:** 10.1101/2023.02.09.527915

**Authors:** Chen Lesnik, Rachel Kaletsky, Jasmine M. Ashraf, Salman Sohrabi, Vanessa Cota, Titas Sengupta, William Keyes, Shijing Luo, Coleen T. Murphy

## Abstract

Reproductive aging is one of the earliest human aging phenotypes, and mitochondrial dysfunction has been linked to oocyte quality decline. However, it is not known which mitochondrial metabolic processes are critical for oocyte quality maintenance with age. To understand how mitochondrial processes contribute to *C. elegans* oocyte quality, we characterized the mitochondrial proteomes of young and aged wild-type and long-reproductive *daf-2* mutants. Here we show that the mitochondrial proteomic profiles of young wild-type and *daf-2* worms are similar and share upregulation of branched-chain amino acid (BCAA) metabolism pathway enzymes. Reduction of the BCAA catabolism enzyme BCAT-1 shortens reproduction, elevates mitochondrial reactive oxygen species levels, and shifts mitochondrial localization. Moreover, *bcat-1* knockdown decreases oocyte quality in *daf-2* worms and reduces reproductive capability, indicating the role of this pathway in the maintenance of oocyte quality with age. Importantly, oocyte quality deterioration can be delayed, and reproduction can be extended in wild-type animals both by *bcat-1* overexpression and by supplementing with Vitamin B1, a cofactor needed for BCAA metabolism.

## Introduction

The progressive inability to reproduce increases with age, typically starting in women’s mid-30s, due to a decline in oocyte quality that precedes the onset of menopause by about 15 years^1,2^. As the maternal age for first childbirth has risen in recent years^3^, understanding the mechanisms that regulate reproductive aging has become increasingly important. Compared to other tissues, oocytes visibly age early in life^1,4,5^, and the decline in oocyte quality with age limits reproduction^5–9^. The ability to reproduce late in life in is determined by oocyte quality in multiple organisms^10–12^, including humans, but the factors that regulate this quality decline with age, and whether oocytes can be rejuvenated in older animals, remain unclear. As in humans, the model organism *C. elegans* undergoes reproductive aging that is regulated by highly conserved genes and pathways that maintain oocyte quality with age^9^. Despite some dissimilarities in reproductive chronologies and strategies between *C. elegans* and humans, the cellular and molecular components that regulate reproductive aging are surprisingly well conserved from worms to humans^9^. Both women and *C. elegans* have long post-reproductive life spans, and both undergo significant reproductive aging on proportional time scales^13^. Most importantly, as in humans, oocyte quality is the critical factor for reproductive aging in worms. We and others have ruled out the possibility that reproductive cessation is limited by oocyte number^8^, germ line stem cells^9^, or apoptosis^9^. Instead, oocyte quality maintenance with age is the critical factor regulating reproductive aging in both *C. elegans* and humans^5,8,9^. We previously found that genes associated with chromosome segregation, cell cycle regulation, DNA damage, and mitochondria decline in aged *C. elegans* oocytes – as in humans - and are critical for oocyte quality maintenance^9^. These genes are higher in young oocytes and in the oocytes of TGF-β mutants, which display slowed reproductive aging^9^. Many of these genes also change with age in mouse^10,12^ and human oocytes^11^, underscoring the similarities in oocyte aging across species.

The insulin/IGF-1 receptor (IIS) mutant *daf-2* provides a compelling model for identifying regulators of healthy aging, as *daf-2* mutations extend lifespan^14^ and generally improve healthspan^15,16^. The IIS pathway also plays a Exploiting *C. elegans’* conserved features of reproductive aging, we addressed these questions through a comparative proteomic analysis. Proteomic analysis of wild-type *C. elegans* revealed age-related changes in protein abundance^28^, but which mitochondrial proteins regulate reproductive longevity are unknown. Here we profiled the mitochondrial proteome of aged wild-type animals and *daf-2* mutants, identifying “youthful” mitochondria-related proteins. We found that the branched-chain amino acid (BCAA) degradation pathway regulates reproductive aging: the first enzyme of the pathway, BCAT-1, is important significant role in regulating the rate of reproductive decline in *C. elegans*^9,13,17^ by increasing oocyte quality^9,18^. IIS and TGF-β pathways are evolutionarily conserved from worms to humans, and are likely to be important in the regulation of human reproductive decline^9,18^. The similarities in reproductive decline between humans and *C. elegans* make worms an excellent model for identifying mechanisms to improve reproductive longevity^19^.

Mitochondria are highly abundant in oocytes, and may play a pivotal role in oocyte aging^20–22^. As human oocytes age, mitochondria activity and number decline, resulting in decreased oocyte quality^5^. However, it is not known whether dysfunction of mitochondrial metabolic pathways - rather than changes in number or morphology^1,5,23–25^-directly regulate oocyte and reproductive aging, or merely correlate with age-related decline. Identifying mitochondrial proteins that change with age may reveal mechanisms that regulate oocyte quality maintenance. Excitingly, interventions aimed at altering aspects of mitochondrial function offer the possibility of slowing various aspects of aging^23,26,27^.for maintaining normal characteristics of mitochondria in *daf-2* oocytes with age. Conversely, increasing BCAA pathway activity in wild-type worms improves oocyte quality and extends reproduction: both overexpression of *bcat-1* and supplementation with Vitamin B1/thiamine, a cofactor needed for the BCAA pathway, extend late-mated reproduction. Our data link mitochondrial BCAA metabolism with oocyte quality maintenance and suggest that modulation of this pathway via Vitamin B1/thiamine may be a strategy to promote reproductive health and longevity.

## Results

### Aged *daf-2* worms maintain ‘youthful’ mitochondria

To identify a set of “youthful” mitochondria-associated proteins associated with *daf-2*’s extended reproductive span (Fig. 1a), we isolated mitochondria using biochemical fractionation and sequential centrifugation from young (Day 1 adult) and reproductively aged (Day 5 adult) wild-type worms and *daf-2(e1370)* mutants (Extended Data Fig. 1). Day 5 of adulthood marks the start of oocyte quality decline in wild-type worms, while both reproduction and high oocyte quality are maintained in *daf-2* animals. LC-MS/MS proteomic analysis identified 1469 proteins with a minimum of two peptides per protein and replication across a minimum of two out of three replicates (Table S1). Mitochondrial proteins from young wild-type and *daf-2* worms are most similar, while aged wild-type mitochondrial proteins differ (Fig. 1b), suggesting that *daf-*2’s mitochondrial proteomes undergo fewer changes in this period. Proteins that are significantly more abundant in the “*daf-2* and young wild-type mitochondria” cluster than in aged wild-type mitochondria may be indicators of mitochondrial functions important for maintaining cellular properties or inhibiting age-related processes. While many proteins are more abundant in old than young worms (Fig. 1c, d), we are interested in proteins that might be associated with improved health; therefore, we focused on proteins that are more abundant in young and *daf-2* animals (Fig. 1e, Table S1). The top “young” mitochondrial proteins (hypergeometric *p-val*< 0.005, Fig. 1e) include BCAT-1 (branched chain aminotransferase), ACDH-1 (acyl CoA dehydrogenase/*dod-12*), W10C8.5 (creatine kinase B), PCYT-1 (phosphocholine cytidylyltransferase), and PHYH-1 (phytanoyl-CoA hydroxylase) (Fig. 1e). Notably, two of these proteins are enzymes in the branched-chain amino acid (BCAA) degradation pathway, suggesting that this pathway could be important for maintaining mitochondrial function with age: BCAT-1, an ortholog of human branched chain amino acid transaminase BCAT1 and BCAT2, is localized to mitochondria in *C. elegans*^29^, and ACDH-1 is an ortholog of human ACADSB (acyl-CoA dehydrogenase short/branched chain).

**Figure 1:**
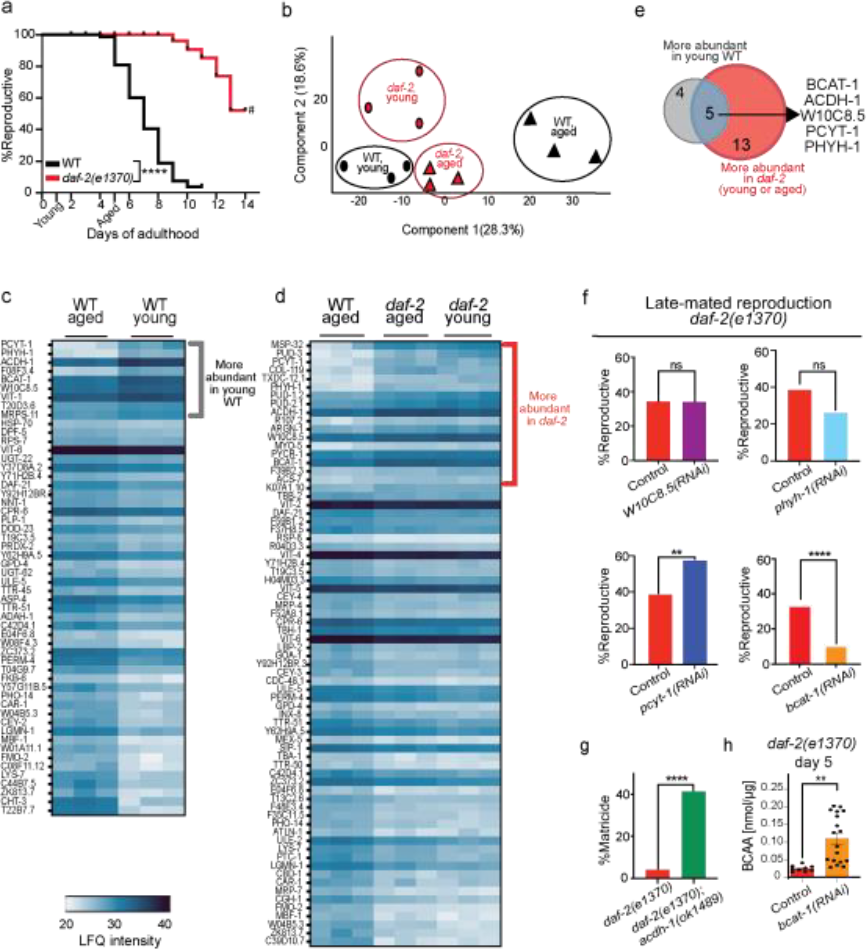
The mitochondrial proteome of *daf-2* is similar to young wild-type and distinct from aged wild-type. (a) *daf-2* mutants have an extended mated reproductive span. Time points of young (Day 1) and aged (Day 5) mitochondria isolation are indicated. # indicates a high matricide frequency. n = 100 worms per group. Mantel-Cox log-ranking, ****p ≤ 0.0001. 3 biological replicates were performed (b) Principal component analysis of mitochondrial proteins suggests a similarity between young wild type and young and aged *daf-2*, and distinct clustering from the aged wild-type mitochondria proteome. (c) LFQ intensity heat maps of proteins that are significantly different (*q-*val< 0.05, -1<log_2_ratio<1) in young wild type (c) or young and aged *daf-2* (d) compared to aged wild-type mitochondria. (e) Venn diagram of proteins that were more abundant in mitochondria isolated from young wild-type and *daf-2* worms compared to aged wild-type worms (5 proteins, hypergeometric *p* < 0.005). The intersection shows the list of shared proteins. (f) Late-mated reproduction of adult Day 8 *daf-2. p-values* (Cochran-Mantel-Haenszel test for multiple chi-square tests, two-sided): *W10C8*.*5* = 0.189, *phyh-1* = 0.12, *pcyt-1* = 0.0039, *bcat-1* = 2.80E^-8^. Control or gene-specific RNAi for *W10C8*.*5* (n = 81, 76), *phyh-1* (n = 96, 92), *pcyt-1* (n = 96, 96), or *bcat-1* (n = 104, 103). (g) *acdh-1* knockout in *daf-2* animals results in high matricide frequency at day 7 of adulthood. n= 74 for *daf-2(e1370)*, n= 68 for *daf-2(e1370);acdh-1(ok1489)*. Chi-square test, two-sided, \*\*\*\**p* ≤ 0.0001. (a, f, g) Representative graph from at least 3 biological repeats. (h) BCAA levels are increased in Day 5 *daf-2* worms after adult-only *bcat-1* knockdown. Each dot represents a different measurement of BCAA concentration from a lysate of at least 70 worms. Control n = 9, *bcat-1(RNAi)* n = 18. Two-tailed, unpaired t-test. \*\**p* = 0.001. Mean ± SEM. All measurements from a total of 3 biological replicates are shown.

The *C. elegans* germline contains the majority of mtDNA in the adult worm^30^, consistent with the energy-intensive role of the germline in reproduction^30–32^. To determine whether the top five ‘youthful’ mitochondrial proteins identified from our proteomic analysis are important for maintaining *daf-2*’s reproduction with age, we tested whether reduction of these genes reduces the late-mated reproduction of aged *daf-2* worms (Fig. 1f). Knockdown of three candidates had either no effect (*W10C8*.*5* and *phyh-1*) or modestly increased reproductive ability (*pcyt-1, p*-val= 0.0039). (Loss of *acdh-1* in *daf-2* animals results in such a high matricide frequency that reproductive span could not be assessed (Fig. 1g).) However, adult-only knockdown of *bcat-1* significantly reduced *daf-2*’s reproductive ability with age (Fig. 1f) and increased BCAA levels (Fig. 1h). Therefore, BCAA metabolism appears to be important for the reproductive success of late-mated *daf-2* animals.

### *bcat-1* is necessary for *daf-2*’s improved reproductive aging

To determine the role of BCAA metabolism genes in *daf-2*’s youthful characteristics, we first tested their effects on *daf-2* lifespan. ACDH-1 acts downstream of BCAT-1, and we previously found that *acdh-1* is required for *daf-2*’s extended lifespan^33^. *bcat-1* reduction also significantly shortens *daf-2*’s lifespan (Fig. 2a, Table S2), suggesting that BCAA metabolism is important for *daf-2*’s longevity.

**Figure 2:**
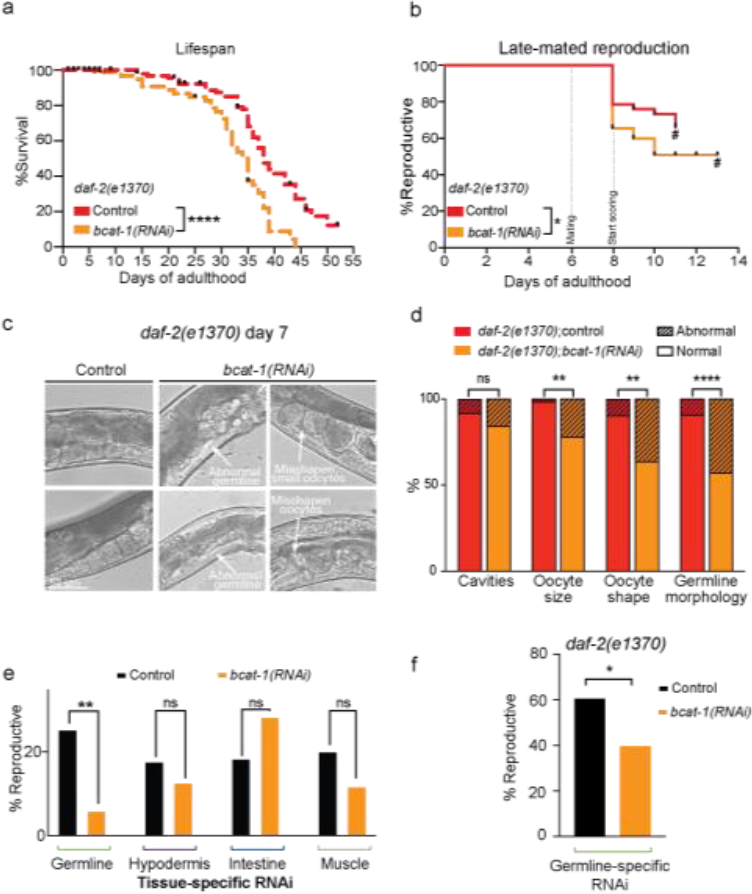
Downregulating *bcat-1* in *daf-2* reduces lifespan, reproductive capability, and oocyte quality. (a) Adult-only knockdown of *bcat-1* shortens the lifespan of *daf-2*. Representative graph. Lifespan data are plotted as Kaplan-Meier survival curves, and the *p-*value was calculated using Mantel-Cox log-ranking (\*\*\*\**p* ≤ 0.0001). n = 120 for each condition. (b) Adult-only knockdown of *bcat-1* decreases the late-mated reproduction of adult *daf-2* at Day 8 as scored every day after mating (n = 75 for control, n= 58 for *bcat-1(RNAi)*, Mantel-Cox log-ranking \**p* = 0.025). (c,d) *bcat-1* is required for oocyte and germline quality maintenance in *daf-2* animals (c, representative images), as scored based on oocyte quality criteria (d). N = 67 for control, n = 57 for *bcat-1(RNAi)*. Chi-square test, two-sided. ns = not significant, \*\**p* = (size = 0.002, shape = 0.004), \*\*\*\**p* ≤ 0.0001. (e) Adult-only knockdown of *bcat-1* specifically in the germline, but not in the hypodermis, intestine, or muscle, decreases the late-mated reproductive capability of Day 5 wild-type worms. *bcat-1 RNAi* was initiated at the L4 stage to avoid developmental lethality and to limit knockdown to adults and the germline/oocytes. Germline: n = 83 for control, n = 70 for *bcat-1(RNAi)*, hypodermis: n = 74 for control, n = 80 for *bcat-1(RNAi)*, intestine: n = 71 for control, n= 74 for *bcat-1(RNAi)*, muscle: n = 45 for control, n = 43 for *bcat-1(RNAi)*, Chi-square test, two-sided. ns = not significant, \*\**p* = 0.0011. (f) Germline-specific *bcat-1* knockdown (adult-only) in *daf-2* animals decreases reproductive ability with age. Late mating was tested at Day 8. N = 82 for control, n = 87 for *bcat-1(RNAi)*, Fisher’s exact test, two-sided, \**p* = 0.014. (a-f) A representative data are shown from (a-e) 3 or (f) 2 biological replicates.

Because *bcat-1* reduction significantly reduced *daf-2*’s late mated reproduction (Fig. 1f), we further investigated the effect of *bcat-1* knockdown; loss of *bcat-1* significantly impairs the ability of *daf-2* mothers to produce progeny after mating (Fig. 2b). Previously, we found that *daf-2’*s ability to reproduce late in life is due to improved oocyte quality maintenance with age^9,18^.

While oocyte quality decline is observed in wild-type worms as early as Day 5, *daf-2* animals maintain high-quality oocytes late into the reproductive period, which extends reproduction^9^. On Day 7 of adulthood, mated *daf-2* animals maintain uniformly sized, cuboidal-shaped oocytes, aligned one next to each other in the proximal end of the gonad^9^ (Fig. 2c, control), but *bcat-1* RNAi treatment causes misshapen and small oocytes and significant germline abnormalities (Fig. 2c, d), suggesting BCAT-1 is required for *daf-2*’s ability to extend reproductive span by maintaining high-quality oocytes with age.

To determine in which tissue BCAT-1 acts to regulate reproductive span, we treated tissue-specific RNAi strains with *bcat-1* RNAi. Knockdown of *bcat-1* in the hypodermis, intestine, or muscle had no significant effect, but reduction of *bcat-1* specifically in the germline significantly reduced the fraction of animals that were reproductive (Fig. 2e). We next tested effect of *bcat-1* reduction specifically in the germline of *daf-2* animals. Similar to wild-type animals, *bcat-1* knockdown specifically in the germline reduced *daf-2*’s reproductive ability with age (Fig. 2f). These results suggest that *bcat-1* acts cell autonomously to regulate reproductive aging.

### *bcat-1*-dependent *daf-2* gene expression in aged oocytes

Because *bcat-1* acts in the germline to regulate reproductive span of *daf-2* worms, differences between *daf-2* and *daf-2;bcat-1(RNAi)* oocytes might reveal oocyte quality-control mechanisms. We carried out RNA sequencing on isolated *daf-2* and *daf-2;bcat-1(RNAi)* day 8 oocytes. Principal component (Fig. 3a) and hierarchical clustering (Fig. 3b) analyses suggest that old *daf-2;bcat-1(RNAi)* oocytes are transcriptionally distinct from old *daf-2* oocytes (*p*-*adj*<0.05, Extended Data Fig. 2, Table S3). *bcat-1* RNAi treatment of *daf-2* worms downregulated many oocyte genes and pathways related to germline and oocyte quality. For example, nine of the 21 *C. elegans skr* (SKp1 Related ubiquitin ligase complex component) genes declined in *daf-2;bcat-1(RNAi)* oocytes (KEGG enrichment (ubiquitin-mediated proteolysis, *padj* = 0.003; Fig. 3C)). Skr/SKP1 genes encode proteins that are in SCF complexes required in mammals for oocyte maturation and embryo development, and their inhibition leads to fertility defects in mouse^34^ and bovine embryos^35^. Delta/lag-2-like Notch ligands were also downregulated in *daf-2;bcat-1(RNAi)* oocytes; GLP-1/Notch signaling regulates functional oocyte development in *C. elegans*^36^, and disruption of Notch signaling in mammals leads to various ovarian pathologies^37^.

**Figure 3:**
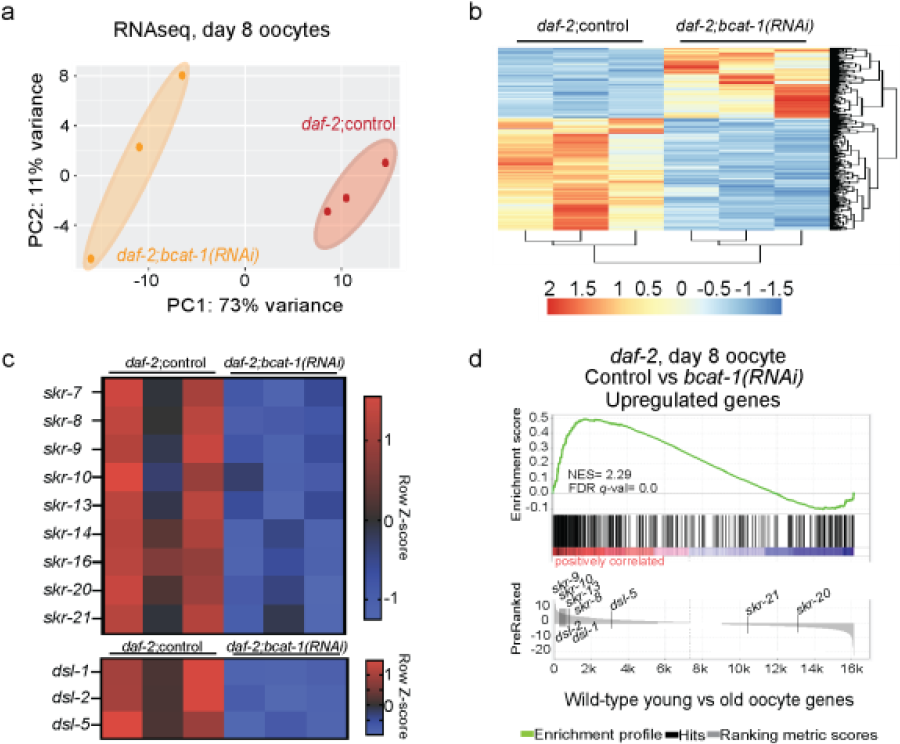
*bcat-1* knockdown alters the expression of oocyte genes important for maintaining youthful characteristics in *daf-2* animals. (a,b) Principal component analysis (PCA) plot (a) and hierarchical clustering (b) of RNA-seq samples from Day 8 oocytes collected from *daf-2*;control and *daf-2;bcat-1(RNAi)* animals. For hierarchical clustering, row Z-scores were determined from normalized counts of differentially expressed genes (DESeq2, padj < 0.05). (c) Heat map plots: Row Z-scores were determined from normalized counts of differentially expressed *skr* genes (SKp1 Related) and dsl genes (Delta/Serrate/Lag-2 domain) from DEseq2. (d) GSEA shows that genes upregulated in *daf-2*;control vs *daf-2;bcat-1* oocytes are enriched in the wild-type young oocytes (Day 1 vs Day 5) gene set. A heatmap of gene clustering from *bcat-1*-dependent *daf-2* genes is displayed on the x axis, while gene enrichment scores compared to young vs old oocyte genes are shown on the y axis. See also Table S3.

We then compared the *bcat-1*-dependent *daf-2* oocyte genes to those that change with age in wild-type oocytes. Gene set enrichment analysis showed that genes upregulated in young oocytes (Day 1 compared to Day 5) are significantly enriched for genes that are more highly expressed in *daf-2* oocytes compared to oocytes from *daf-2;bcat-1(RNAi)* worms (Fig. 3d). These results suggest that *bcat-1* is required by *daf-2* animals, at least in part, to maintain the youthful gene expression characteristics of its oocytes.

### *bcat-1* loss alters mitochondria localization and activity

We next examined how *bcat-1* reduction affects germline mitochondrial function and morphology. To visualize germline mitochondria, we used a *daf-*2 strain expressing the outer mitochondrial membrane protein TOMM-20 fused to mKate2 specifically in the germline. Under control RNAi conditions, mitochondria are uniformly distributed in mature oocytes (Fig. 4a, control). However, adult-only knockdown of *bcat-1* in *daf-2* induces perinuclear clustering of oocyte mitochondria (Fig. 4a, b, inset, Supp. Videos 1-3). Perinuclear mitochondrial distribution is associated with cellular stress and changes in cellular ROS (reactive oxygen species) levels^38^, and we previously found that *bcat-1* knockdown increases mitochondrial hyperactivity, increasing the mitochondrial oxygen consumption rate^39^.

**Figure 4:**
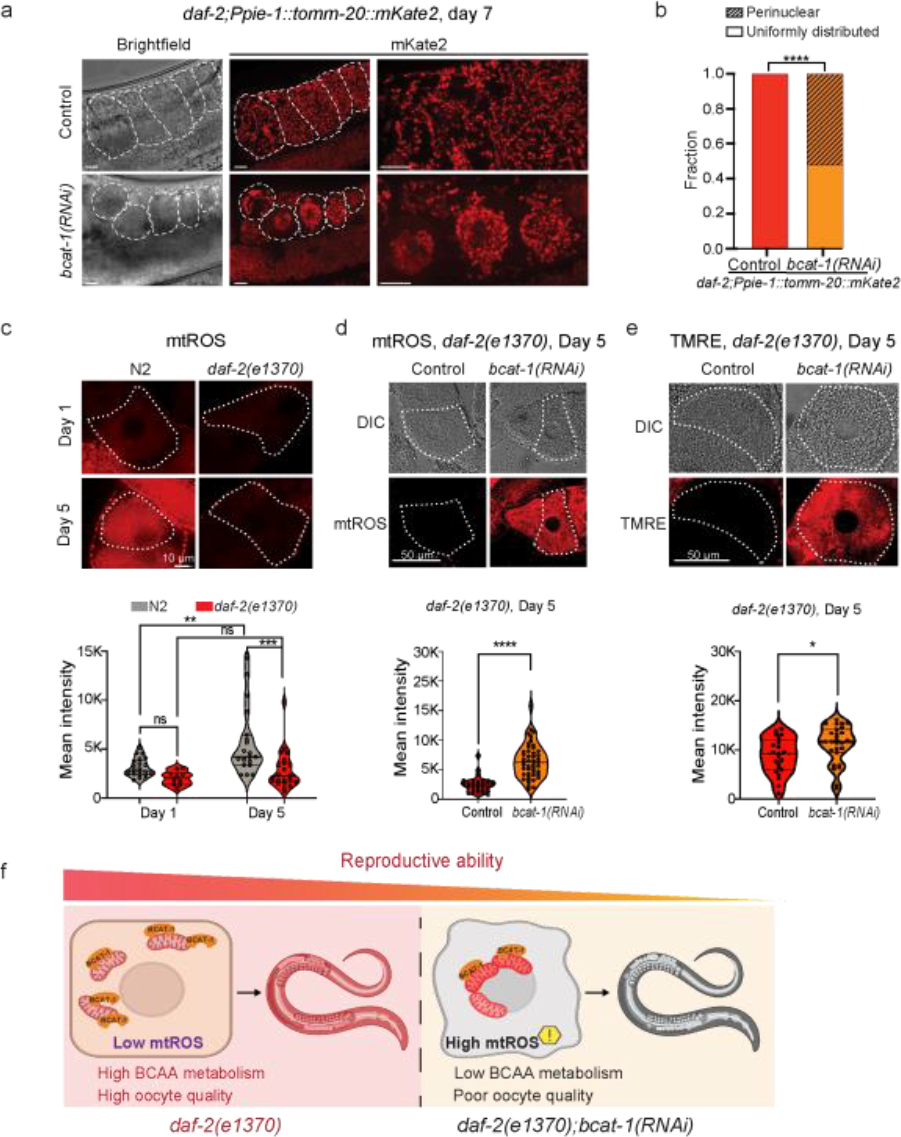
*bcat-1* knockdown in *daf-2* mutants alters mitochondria cellular localization, mtROS levels, and mitochondria activity. (a) Adult-only *bcat-1* knockdown results in the perinuclear localization of mitochondria in *daf-2* oocytes on day 7. A white dashed outline marks mature oocytes. Scale bars: 10 μm (b) Quantification of perinuclear mitochondria localization in *daf-2* oocytes from (a). Representative graph from 3 biological replicates. N = 36 for control, n = 44 for *bcat-1(RNAi)*. Chi-square test, two-sided. \*\*\*\**p* ≤ 0.0001. (c) mtROS levels are increased on day 5 in the mature oocyte of wild-type animals, but not *daf-2* (compared to day 1). Bottom: quantification of mtROS (mean intensity) in -1 oocyte in wild-type animals compared to *daf-2*, Day 1 vs. Day 5 of adulthood. Day 1 n, (N2 = 22, *daf-2* = 10), Day 5 n, (N2 = 19, *daf-2* = 22). Two-way ANOVA with Tukey’s multiple comparison. ns = not significant, \*\**p* = 0.0018, ***p = 0.0005. (d) mtROS levels are increased in *daf-2* mature oocytes on Day 5 upon adult-only *bcat-1* knockdown. Bottom: mtROS levels in *daf-*2 mature oocytes on day 5 upon adult-only *bcat-1* knockdown. N = 30 for control, n = 46 for *bcat-1 (RNAi)*. Two-tailed, unpaired t-test. \*\*\*\**p* ≤ 0.0001. (e) Mitochondrial membrane potential is higher in *daf-*2 mature oocytes on Day 5 upon adult-only *bcat-1* knockdown. Bottom: quantification of TMRE signal in *daf-2* mature oocytes on Day 5 upon adult-only *bcat-1* knockdown. Control n = 23, *bcat-1(RNAi)* n = 27. Two-tailed, unpaired t-test. \**p* = 0.034. For (c-e), a white dashed outline marks the most mature oocyte (−1 oocyte). (a-e) Representative images and graphs from 3 biological replicates are shown. (f) Model: *bcat-1* is important for mitochondria cellular localization, mtROS levels, and mitochondria activity in *daf-2* mature oocytes. Upon *bcat-1* downregulation, mitochondria are altered and oocyte quality is poor, resulting in a shorter reproductive span. Created with BioRender.com.

Therefore, we tested whether BCAA metabolism affects mtROS in aged *daf-*2 oocytes using the MitoTracker Red CM-H2XRos reporter (Extended Data Fig. 3a). Unlike wild-type oocytes, which show increased mtROS levels with age, *daf-2* maintains low mtROS in mature oocytes with age (Fig. 4c). However, mtROS levels are higher in Day 5 *daf-2;bcat-1(RNAi)* oocytes compared to the control (Fig. 4d), suggesting that *bcat-1* is essential for maintaining low levels of cellular mitochondrial ROS. Maintaining mitochondria membrane potential (MMP), which in turn maintains appropriate ROS levels, is crucial for preserving mitochondrial health and ensuring overall cellular function. To specifically assess MMP, we used tetramethylrhodamine ethyl ester (TMRE), a dye that labels active mitochondria through increased mitochondrial membrane potential (Extended Data Fig. 4). *bcat-1* knockdown increased TMRE fluorescence intensity in *daf-2* oocytes, indicating mitochondrial hyperactivity (Fig. 4e), consistent with our previous results^39^. Unlike the effects of *bcat-1* knockdown, we found no significant differences in mtROS levels (Extended Data Fig. 4a) and MMP (Extended Data Fig. 4b) in mature oocytes upon *acdh-1* knockdown in aged *daf-2* animals. Similarly, we found no significant differences in mtROS levels comparing aged *acdh-1* vs. wild-type oocytes (Extended Fig. 4c). In contrast to BCAT-1, which is essential for the metabolism of all three BCAAs, ACDH-1 is specialized for the processing of isoleucine and valine. These results suggest that the specific metabolites upstream of or in parallel to *acdh-1* regulate mitochondrial function in aged *daf-2* oocytes.

Together, these results indicate that altered BCAA metabolism caused by *bcat-1* reduction disrupts mitochondrial cellular distribution and promotes hyperactive mitochondria in *daf-2* oocytes. *bcat-1* reduction is also associated with elevated mtROS levels in aged oocytes, like those observed in wild-type oocytes with age. These changes upon reduction of BCAT-1 may contribute to the accelerated decline of *daf-2* oocyte quality and reduction of reproductive span (Fig. 4f).

### *bcat-1* overexpression improves reproduction oocyte quality

We next asked how changes in *bcat-1* levels affect wild-type animals. As in *daf-2* mutants (Fig. 1h), reduction of *bcat-1* increases BCAA levels in wild-type worms (Fig. 5a). Both late-mated reproduction (Extended Data Fig. 5a) and reproductive span (Fig. 5b, Table S2) of wild-type worms are reduced upon *bcat-1* knockdown, demonstrating that *bcat-1* is required for normal reproduction. To test whether increasing BCAA levels individually would mimic *bcat-1* knockdown’s effect on reproduction, we supplemented to wild-type adult worms with each BCAA individually (5 mM in plate) and monitored reproductive span. Of the three BCAAs, supplementing leucine alone was sufficient to shorten the reproductive span, but the magnitude is smaller than *bcat-1* knockdown (Fig. 5c).

**Figure 5:**
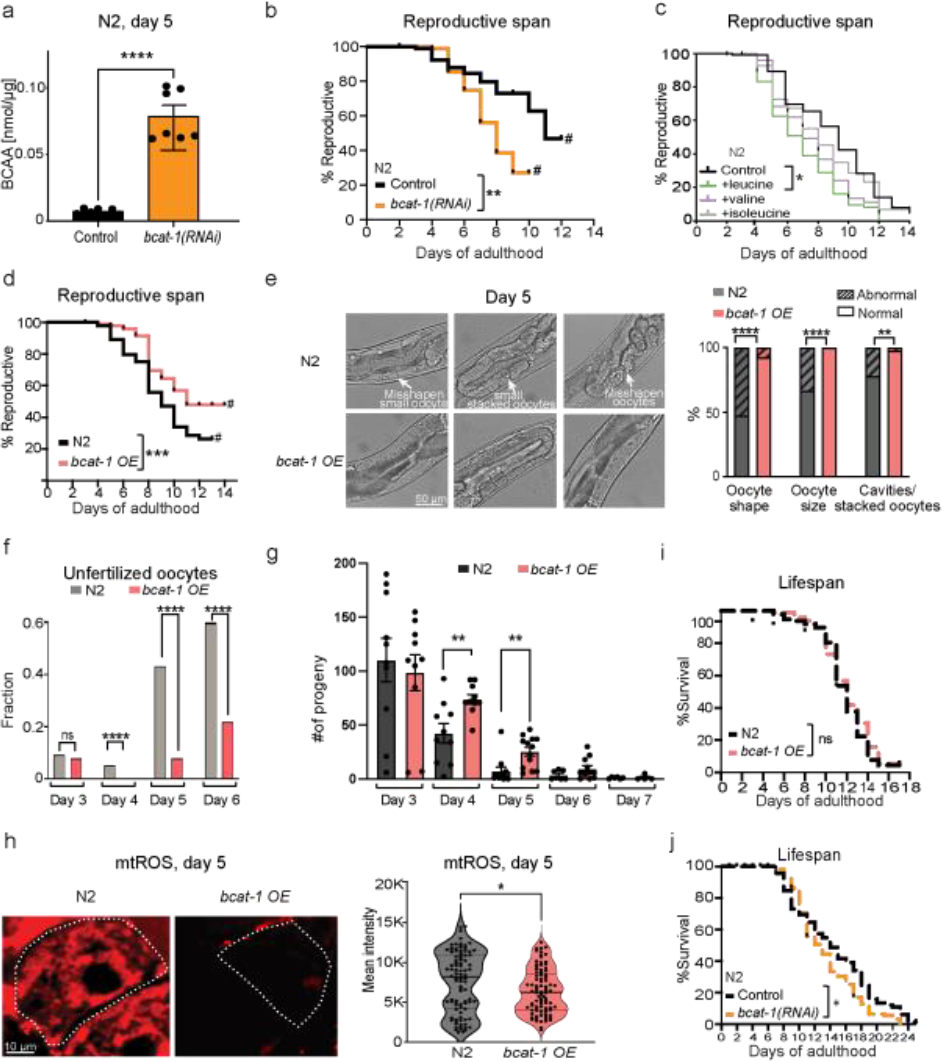
*bcat-1* knockdown reduces the reproductive capability of wild-type worms, while its overexpression improves wild-type oocyte quality and extends reproductive span. (a) BCAA levels increase after adult-only *bcat-1* knockdown. Each dot represents BCAA concentration measurements from lysate containing at least 70 worms. 3 biological replicates are shown. Two-tailed, unpaired t-test. \*\*\*\**p* ≤ 0.0001. Mean ± SEM. (b) *bcat-1* knockdown reduces mated reproductive span of wild-type worms. n = 100 for control, n = 90 for *bcat-1(RNAi)*. \*\**p* = 0.0016. (c) 5 mM leucine shortens mated wild-type reproductive span. N = 100 for each condition. \**p* = 0.04 (d) Overexpressing *bcat-1* (*bcat-1 OE*) extends mated reproductive span. Wild type (n = 100), *bcat-1* OE (n = 90), \*\*\**p* = 0.0008. (e) *bcat-1* overexpression improves oocyte quality with age. Wild type (n = 36), *bcat-1* OE (n = 40). \*\*\*\**p* ≤ 0.0001, \*\**p* = 0.008. (f) Compared to wild type, mated *bcat-1* OE produce fewer unfertilized oocytes (relative to fertilized embryos). Chi-square tests. ns = not significant, \*\*\*\**p* ≤ 0.0001. (g) Progeny production in wild type compared to *bcat-1 OE*. Unpaired t-test. \*\**p* (Day 4 = 0.006, Day 5 = 0.004). Wild type or *bcat-1 OE* animals analyzed on Day 3 (n = 10, 10) Day 4 (n = 10, 10), Day 5 (11, 13), Day 6 (7, 11), or Day 7 (n = 6, 4). Mean ± SEM. (h) Left: mtROS levels are lower in the mature oocyte of aged *bcat-1* OE worms compared to wild type. White outline marks the most mature (−1) oocyte. Right: mtROS quantification (mean intensity) in the -1 oocyte of Day 5 wild-type compared to *bcat-1 OE*. Data pooled from three biological replicates. Unpaired t-test. N2 (n = 94), *bcat-1OE* (n = 89), \**p* = 0.018. (I, j) *bcat-1* knockdown (i) and overexpression (j) lifespans. (i) n = 100, and (j) n = 120 for each condition. Data plotted as Kaplan-Meier survival curves, Mantel-Cox log ranking (ns = not significant, *p = 0.033). (a-j) Representative data from at least 3 biological replicates are shown.

We next asked whether increasing BCAT-1 levels could slow reproductive decline, using a published and verified *bcat-1* overexpression strain^40^ (Extended Data Fig. 5b). Indeed, overexpression of *bcat-1* significantly extends reproductive span (Fig. 5d, Table S2). To determine whether *bcat-1* overexpression slows oocyte quality loss with age, we examined oocytes on Day 5 of adulthood following mating; *bcat-1* overexpression significantly improved oocyte quality relative to wild-type controls (Fig. 5e). Like human oocytes, *C. elegans* oocytes similarly display age-related oocyte developmental defects, such as chromosomal segregation errors that result in reduced oocyte fertilizability and embryo hatching^9,41^. We previously showed that the oocytes of aged mated wild-type hermaphrodites have reduced chromosome segregation fidelity and that these animals produce significantly more unfertilized oocytes compared to young worms^9^. To determine whether *bcat-1* overexpression improves oocyte fertilizability with age, we mated wild-type hermaphrodites and counted the proportion of unfertilized oocytes as the animals aged. Consistent with our previous results^9^, wild-type animals produced a significantly greater proportion of unfertilized oocytes with age. However, worms overexpressing *bcat-1* produced significantly fewer unfertilized oocytes relative to fertilized embryos (Fig. 5f). *bcat-1* overexpression also may increase progeny production with age (Fig. 5g).

Wild-type oocytes exhibit increased levels of mtROS with age (Fig. 4c, d), while ROS levels are decreased in *daf-2* oocytes, and Day 5 *daf-2;bcat-1(RNAi)* oocytes show higher mtROS levels compared to the *daf-2;control(RNAi)* group (Fig. 4e, f), implying a crucial role for *bcat-1* in maintaining low levels of cellular mtROS. Consistent with these results, BCAT-1 overexpression leads to reduced mtROS levels in aged wild-type oocytes (Fig. 5h).

Since *bcat-1* overexpression is beneficial for reproductive aging, we next examined the role of *bcat-1* in lifespan (Fig. 5i, j; Table S2). We found that neither overexpression nor knockdown of *bcat-1* affected lifespan (Fig. 5i, j, Extended Data Fig. 5c, Table S2), counter to a previous report^40^. While we observed effects on other age-related phenotypes upon *bcat-1* knockdown, including decreased reproductive span, increased BCAA levels (Fig. 5a), and changes in mitochondrial ROS, we never observed an extension of lifespan (Extended Data Fig. 5c) even using the same reagents^40^. Thus, *bcat-1* levels appear to regulate oocyte quality and reproductive aging but have no apparent role in the regulation of lifespan in wild-type animals. These findings parallel our prior study demonstrating the potential decoupling of extended reproductive span and lifespan^13^. Overall, these results suggest that increased levels of BCAT-1 are sufficient to improve reproductive aging and oocyte quality in *C. elegans*.

### BCAT-1 overexpression and *daf-2* share oocyte transcripts

To identify oocyte transcriptional changes that might regulate oocyte quality maintenance, we performed RNA sequencing on unfertilized oocytes isolated from *fem-1* (self-spermless) control and *fem-1*;*bcat-1*-overexpressing worms on day 8 of adulthood, and compared the suite of significantly changed *bcat-1* overexpression-regulated oocyte genes (*padj*<0.05, Extended Data Fig. 6, Table S4) to previously identified *daf-2(RNAi)* oocyte transcriptional targets^18^. Genes upregulated in aged *daf-2(RNAi)* oocytes are highly enriched among the upregulated genes in *bcat-1*-overexpressing worms’ oocytes (Fig. 6c). Similarly, downregulated *daf-2(RNAi)* oocyte targets are enriched for genes that decline in oocytes from *bcat-1*-overexpressing worms (Fig. 6d). These findings suggest that *bcat-1* overexpression regulates oocyte gene expression through a shared subset of genes also targeted by the *daf-2*/IIS pathway. Several of the oocyte transcriptional targets upregulated by *bcat-1* overexpression and *daf-2(RNAi)* were previously shown to have functional roles in the germline and reproduction. For example, *scl-19* encodes an SCP-like secreted protein that affects germline tumor formation^42^. Oocytes from both *daf-2(RNAi)* and *bcat-1*-overexpressing worms had significantly upregulated *folt-1* levels, and *folt-1* (folate transporter) mutants exhibit many germline and oocyte defects^43^. These findings are consistent with studies showing that the folate receptor is expressed in oocytes and that maternally contributed folate receptor transcripts promote embryonic development^44^.

**Figure 6:**
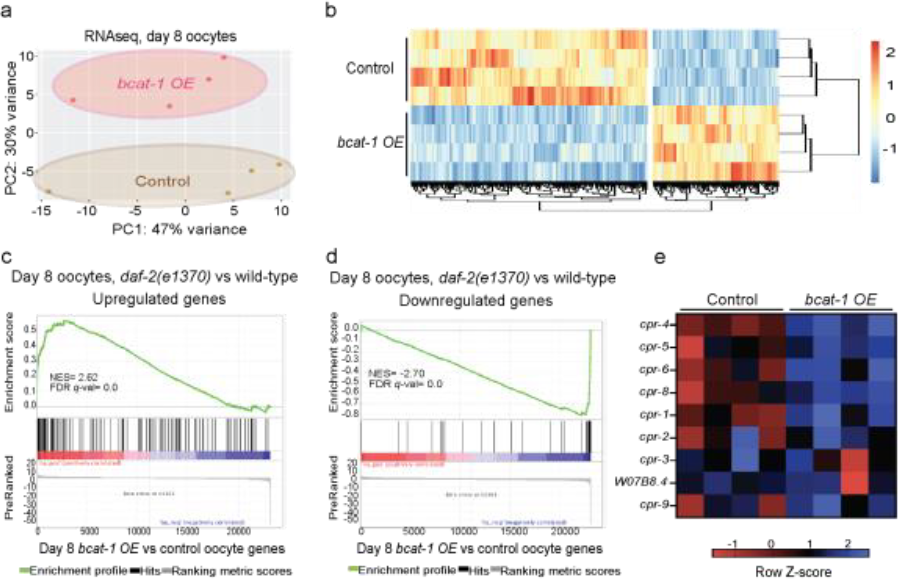
Oocytes from *bcat-1*-overexpressing are transcriptionally similar to oocytes from *daf-2(RNAi)* worms. (a) Principal component analysis (PCA) plot of RNA-seq samples from Day 8 oocytes collected from *fem-1* control and *fem-1;bcat-1* overexpressing animals. (b) Row Z-scores were determined from normalized counts of differentially expressed genes (DESeq2, padj < 0.05), and hierarchical clustering was performed. (c,d) GSEA shows that genes (c) upregulated in Day 8 *daf-2* oocytes^18^ are enriched in the upregulated genes from the *bcat-1* overexpression vs control gene set, while (d) downregulated Day 8 *daf-2* oocytes are enriched in the downregulated genes in oocytes from *bcat-1*-overexpressing worms (e) *bcat-1*-overexpressing worms downregulate cathepsin protease genes in oocytes. Row Z-scores were determined from normalized counts of *C. elegans cpr* genes (cysteine protease related) from DEseq2. See also Table S4.

Reduction of *daf-2* results in the downregulation of cathepsin-B-like cysteine proteases in aged oocytes^18^, and reduction of cathepsin B activity by RNAi knockdown or by pharmacological cathepsin B enzyme inhibition improved oocyte quality with age in wild-type animals^18^. Similarly, significant downregulation of several *cpr* (cysteine protease related) genes was observed in oocytes from *bcat-1*-overexpressing worms (Fig. 6e). Additional shared downregulated transcriptional targets include vitellogenins that encode major yolk proteins, and the metallothionein *mtl-2*. Overall, the similarity between oocytes from *daf-2(RNAi)* and *bcat-1*-overexpressing worms suggests that *bcat-1* overexpression acts at least partially through shared *daf-2*/IIS targets to improve reproductive span and oocyte quality with age.

### Vitamin B1 slows reproductive decline in wild-type worms

BCAT-1 catalyzes the first step of the catabolism of branched-chain amino acids (BCAAs) (Fig. 7a), and we found that its reduction is sufficient to elevate BCAA levels in aged wild-type and *daf-2* animals (Fig. 1h, Fig. 5a). Our findings suggest that maintaining BCAT-1 levels with age may extend reproduction by maintaining flux through the BCAA pathway. We hypothesized that other treatments that affect BCAA metabolism downstream of BCAT-1 might similarly improve reproductive aging in *C. elegans*. The rate-limiting reaction of the BCAA metabolism pathway is catalyzed by the branched-chain α-keto acid dehydrogenase (BCKDH) complex^45^ (Fig. 7a). This mitochondrial enzymatic complex converts the α-keto acids of each branched-chain amino acid to the corresponding acyl CoAs. BCKDH is composed of three subunits, E1, E2, and E3. The E1 subunit binds thiamine pyrophosphate (ThPP), which is an essential cofactor and a derivative of thiamine, also known as Vitamin B1^46^. Animals cannot synthesize thiamine and rely on its absorption from their diet. After being imported into animal cells, thiamine is transformed into ThPP by thiamine pyrophosphokinase^46^. When thiamine levels are insufficient, BCAA catabolism is impaired, increasing the levels of unprocessed branched-chain keto acids^47^.

**Figure 7:**
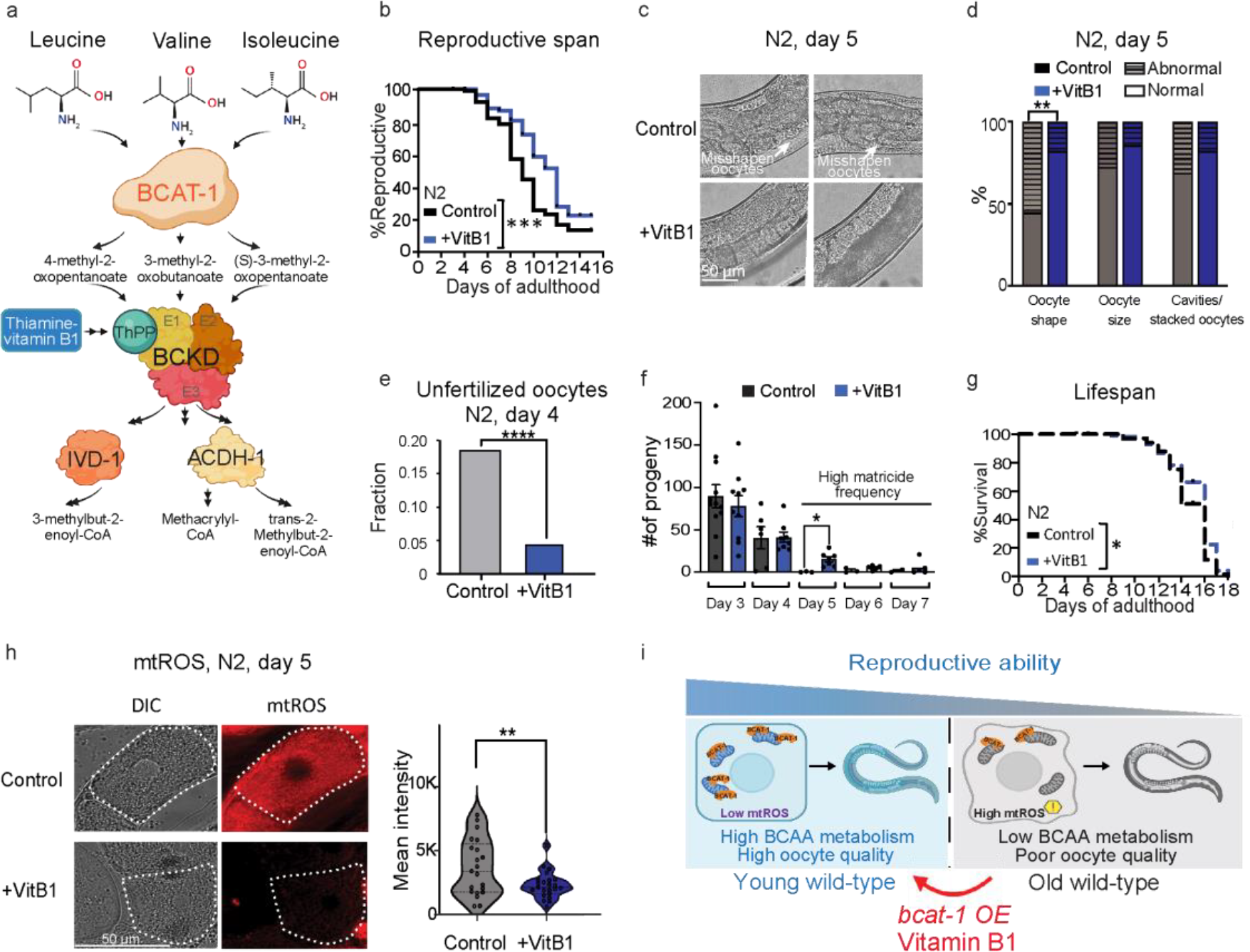
Adult-only Vitamin B1 supplementation improves wild-type oocyte quality and extends reproductive span. (a) Schematic of selected steps in the branched-chain amino acid metabolism pathway in *C. elegans*. (b) Supplementation with 5 mg/mL of thiamine (Vitamin B1) starting on day 1 of adulthood significantly extends the mated reproductive span of wild-type animals. N = 90 for control, n = 88 for Vitamin B1 condition. Representative graph of 3 biological repeats. Mantel-Cox log-ranking \*\*\**p* = 0.0003. (c) Adult-only supplementation of Vitamin B1 is sufficient to maintain better oocyte quality with age. (d) Day 5 oocytes quality quantification in control and Vitamin B1-supplemented worms. Control n = 29, Vitamin B1 n = 28. Chi-square test, two-sided. \*\**p* = 0.0035, ns = not significant. (e) Mated animals supplemented with Vitamin B1 during their adulthood produce fewer unfertilized oocytes (relative to fertilized embryos) compared to control animals. Chi-square test, two-sided. \*\*\*\**p* ≤ 0.0001. Control (embryo n = 261, unfertilized n = 59), or Vitamin B1 (embryo n = 329, unfertilized n = 15). (f) Progeny production in control or Vitamin B1-supplemented animals. Unpaired t-test, two-sided. \**p* = 0.016. Wild type or Vitamin B1-supplemented animals analyzed on Day 3 (n = 10, 10) Day 4 (n = 6, 8), Day 5 (3, 7), Day 6 (3, 5), or Day 7 (n = 2, 5). Mean ± SEM. (g) Supplementing Vitamin B1 starting on day 1 of adulthood slightly extends lifespan (3%, Mantel-Cox log-ranking p = 0.033). (h) Left: mtROS levels were significantly lower in the Day 5 mature oocytes of wild-type worms supplemented with Vitamin B1 compared to controls. A white dashed outline marks the most mature oocyte (−1 oocytes). Representative images. Right: Representative graph of 3 biological repeats. Control n = 21, Vitamin B1 n = 23. Two-tailed, unpaired t-test. \*\**p* = 0.0094. (i) BCAT-1 is important for wild-type reproduction. BCAT-1 levels are reduced with age, resulting in reduced reproduction with age due to declining oocyte quality. By overexpressing *bcat-1* or by supplementing with Vitamin B1 during adulthood, reproduction can be extended. (a, i) Created with BioRender.com. (b-h) Representative data from 3 biological replicates are shown.

To test whether Vitamin B1 affects reproductive aging, we supplemented the bacterial diet with Vitamin B1 starting on the first day of adulthood; this treatment significantly extends the mated reproductive span of wild-type worms, increasing the median reproductive span by more than 30 percent (Fig. 7b, Table S2). Moreover, oocytes of aged animals supplemented with Vitamin B1 were significantly higher in quality, particularly in oocyte shape (Fig. 7c, d). We also found that hermaphrodites supplemented with Vitamin B1 at day 4 of adulthood produced fewer unfertilized oocytes, similar to worms overexpressing *bcat-1* (Fig. 7e) and other mutants with extended reproductive spans (i.e., *daf-2* and *sma-2)*^9^, suggesting that Vitamin B1 supplementation delays not just oocyte deterioration, but also the decline in oocyte fertilizability with age.

We next examined whether Vitamin B1 has similar effects on lifespan compared to its effect on reproductive span. Adult-only supplementation of Vitamin B1 very slightly extends the lifespan of wild-type animals (Fig. 7g, Table S2), but to a lesser extent than its effect on reproductive span (Fig. 7b). These results suggest a specific effect of Vitamin B1 on reproduction, rather than somatic health with age, but also suggest that Vitamin B1 is not toxic to the animals.

Mitochondrial ROS (mtROS) levels are elevated in both aged wild-type and aged *daf-2(e1370);bcat-1(RNAi)* worms (Fig. 4c-f). Because supplementing Vitamin B1 during adulthood is sufficient to improve oocyte quality with age and to extend reproduction, we next tested whether supplementation with Vitamin B1 also reduces mtROS in aged oocytes. Indeed, aged oocytes of animals supplemented with Vitamin B1 have significantly lower levels of mtROS compared to untreated animals (Fig. 7h). Together, these data suggest that the addition of a dietary supplement that promotes BCAA metabolism starting in adulthood is sufficient to improve oocyte quality, thus extending reproductive ability with age at least in part through a reduction in mtROS levels in aged oocytes (Fig. 7i).

## Discussion

Here, we have established a regulatory connection between BCAA metabolic enzymes and reproductive longevity. By profiling the mitochondrial proteome of young and old wild-type and *daf-2* insulin/IGF-1 mutant *C. elegans*, we discovered a “youthful” mitochondrial proteomic profile that is distinct from that of aged animals; the level of the first enzyme in BCAA metabolism, BCAT-1, is one of the principal differences in these profiles.

We found that *bcat-1* is crucial for *daf-2*’s maintenance of reproduction with age through its regulation of mitochondrial localization, mitochondrial activity, and low mtROS levels in *daf-2*’s aged oocytes. Our transcriptional data indicate that *bcat-1* is also required for *daf-2* oocytes to maintain youthful gene expression. Additionally, we found that *bcat-1* overexpression extends wild-type reproductive span by improving oocyte quality. Furthermore, supplementation of the BCAA-pathway co-factor Vitamin B1 improves age-related phenotypes (e.g., oocyte fertilizability, mitochondrial localization and function, and mtROS levels) in oocytes to extend the reproduction of wild-type animals.

We found that reduction of *bcat-1* disrupts BCAA metabolism, increasing levels of BCAT-1 substrates (BCAAs) and reducing the reproductive capability of both wild-type and *daf-2* mutants. Our results are consistent with a recent study of the *bcat-1(hh58)* mutant, which also exhibits elevated BCAA levels and significant reproductive and developmental abnormalities^48^. Unlike *bcat-1*, deletion of *acdh-1*, which encodes the enzyme ACADSB and is specialized for the processing of isoleucine and potentially valine^49,50^ (and is the first enzyme in the alternate propionyl-CoA shunt catabolism pathway^49^), had no significant effect on mitochondrial mtROS; furthermore, we found that supplementation of leucine alone is sufficient to shorten reproductive span, although to a lesser extent than *bcat-1* knockdown. These results suggest that metabolites upstream of or in parallel to *acdh-1* regulate mitochondrial function in aged *daf-2* oocytes. Our study aligns with prior research demonstrating the distinct roles of individual BCAAs in influencing specific phenotypes. For instance, in mouse and human studies, reducing dietary isoleucine or valine, but not leucine, enhances metabolic health in mice, while higher isoleucine intake in humans correlates with increased BMI^51^. In *Drosophila*, a decrease in isoleucine, but not valine, leucine, or their combinations, was found to be crucial for stimulating feeding behavior^52^. These suggest that individual BCAAs have specific influences on various phenotypes.

We previously found that downregulation of neuronal *bcat-1* increases endogenous leucine/isoleucine levels, mitochondrial hyperactivity, ROS damage, and subsequent age-dependent motor dysfunction and degeneration of dopamine neurons with age, phenocopying a Parkinson’s Disease phenotype^39^; these results led us to examine *BCAT1* expression in the substantia nigra of sporadic Parkinson’s Disease patients, which we found was also significantly downregulated^53^. Alzheimer’s Disease (AD) mice also have elevated plasma levels of BCAAs, and *Bcat1* was downregulated in their brain tissues^54^. Taken together, these findings suggest that reduction of *bcat-1* is linked to increased BCAA levels, a disrupted flux of BCAA metabolism, and subsequent deleterious effects on age-related phenotypes, while increased *bcat-1* overexpression reverses these effects. Consistent with our results, BCAA metabolic enzyme genes are upregulated in both the long-lived mutants *daf-2* and *eat-2* mutants compared to wild-type animals^55^; metabolomic profiling of these mutants revealed decreased endogenous BCAA levels, suggesting an increased flux through the BCAA pathway. Similarly, a BCAA-restricted diet extends Drosophila lifespan^56^, restriction of dietary BCAA levels also extends the lifespan of two progeroid mouse models, and a reduced BCAA diet started at midlife improves metabolic health and decreases frailty of a wild-type mice strain^57^. Together, these results suggest that decreased endogenous BCAA levels and increased flux through the BCAA metabolism pathway work similarly to improve age-related characteristics.

We found that overexpression of *bcat-1* or supplementation with Vitamin B1 maintains low oocyte mtROS, promotes oocyte quality, and extends reproductive longevity. Our previous metabolomic analyses showed that the levels of TCA cycle substrates and precursors, such as pyruvate and malate, are significantly decreased in *bcat-1(RNAi)* worms, suggesting a potential link between BCAA and TCA cycle metabolism^39^. Recently, crosstalk between BCAA and pyruvate metabolism was shown: BCKAs inhibit the mitochondrial pyruvate carrier^58^, and mitochondrial pyruvate carrier inhibition stimulates BCAA catabolism^59^. Under normal conditions, BCAT-1 converts BCAAs into their corresponding BCKAs that inhibit pyruvate uptake^58^ and maintain normal ROS levels.

Supporting this model, *acdh-1* knockdown, which likely only increases the level of the BCKA for isoleucine, but not those for leucine or potentially valine^49,50^, neither affects *daf-2’s* mtROS or TMRE levels (Extended Data Fig. 4). Conversely, downregulating *bcat-1* results in accumulation of BCAA substrates (Fig. 1h, 5a), which may decrease BCKAs levels that then stimulate pyruvate uptake^58^.

Mitochondrial pyruvate uptake hyperpolarizes the MMP, resulting in the stimulation of ROS production^60^. Therefore, overexpressing *bcat-1* or supplementing with Vitamin B1, which we found to promote oocyte and reproductive longevity in wild-type animals, may maintain low mtROS levels through an increase in BCAA metabolism. Interestingly, Vitamin B1 activation of BCKA dehydrogenase would likely decrease BCKA levels^61^, which may increase pyruvate metabolism and mtROS production. Consequently, Vitamin B1 may only provide reproductive benefits by decreasing BCAAs without increasing BCKA levels.

The conserved mechanisms of reproductive aging and BCAA metabolism in *C. elegans* and humans provide unique opportunities to identify safe and effective treatments to slow reproductive decline with age. Here we found that both BCAT-1 overexpression and Vitamin B1 supplementation promote oocyte and reproductive health in wild-type animals. Vitamin B1 was previously shown to increase the activity of human liver BCKD^62^, which catalyzes the flux-limiting step of BCAA metabolism. Beyond its well-established role as a cofactor for BCKD, Vitamin B1 also serves as a cofactor for several enzymes involved in various metabolic processes.

While we cannot rule out the influence of Vitamin B1 as a cofactor for other enzymes that could regulate reproductive span, it is unlikely that this is due to Vitamin B1-induced pyruvate dehydrogenase or α-ketoglutarate dehydrogenase activation. If increased pyruvate dehydrogenase or α-ketoglutarate dehydrogenase activity were responsible for the reproductive benefits of Vitamin B1, we would have expected increases in lifespan similar to those previously reported for pyruvate (14%)^63^ or α-ketoglutarate (50%)^64^ supplementation. However, the strikingly similar reproductive phenotypes observed between Vitamin B1 supplementation and *bcat-1* overexpression, including increased reproductive span, oocyte quality, oocyte fertilizability, and lower mtROS, suggest that Vitamin B1 acts in the same pathway as BCAT-1. Vitamin B1 supplementation experiments were performed on heat-killed bacteria to prevent the interfering effects of bacterial metabolism^65^, limiting our ability to directly test the role of BCKA dehydrogenase in Vitamin B1-induced reproductive benefits using RNAi knockdown; nor can we test *bckd* subunits, as these mutants are not fertile^66,67^. We thus hypothesize that increased BCAA metabolism, either through overexpression of *bcat-1* or supplementation with Vitamin B1 to activate BCKD, slows age-related oocyte decline and extends reproductive span. These beneficial effects are unlikely to be due to reversal of Vitamin B1 malnutrition, as we observed no large changes in lifespan with these treatments.

While a possible link between Vitamin B1 and oocyte maturation and quality has been shown in mice^68^, those results were found in the context of Vitamin B1 deficiency in young reproductive females, not in aged wild-type animals, as we studied here. Therefore, the pathology associated with Vitamin B1 deficiency is likely distinct from the beneficial effects that Vitamin B1 supplementation has on improved oocyte quality and reproduction. This is not the only instance where differences in mechanisms and phenotypic outcomes between dietary deficiency and supplementation are apparent, and such distinctions are consistently observed across various species, especially as individuals age. Supplementation of biomolecules – even in the absence of obvious deficiency – can be beneficial. Coenzyme Q10, for example, extends lifespan and reduces oxidative stress in wild-type *C. elegans* under normal growth conditions^69^. No adverse effects have been reported for Vitamin B1/thiamine supplementation^70^, suggesting that this is may be a safe intervention to increase available thiamine levels, and as our study suggests, may increase oocyte quality in females with age.

Branched-chain amino acid metabolism is a highly conserved and essential process from bacteria to mammals^71^. Our study establishes links between mitochondrial BCAA metabolism, mtROS levels, oocyte quality, and reproductive aging. Our data show that age-related decline in mitochondrial BCAT-1 levels correlates with increased oocyte mtROS, and elevated mtROS levels resulting from *bcat-1* knockdown appear to promote oocyte aging. This role of mtROS in regulating oocyte quality in *C. elegans* is supported by the oocyte and reproductive health effects associated with small molecules that induce antioxidant pathways^23,26^. Additionally, mitochondria in both human and *Xenopus* early oocytes have low MMP and ROS levels as a result of the absence of complex I in these oocytes; this lack of ROS production promotes the longevity of oocytes^22^. Similarly, *Drosophila* oocyte mitochondria shift toward a low-activity state at the end of oogenesis, which is required for developmental competence after fertilization^72^. Similar to the effect we observed on mitochondrial morphology in aged *daf-2;bcat-1(RNAi)* oocytes, abnormal mitochondrial aggregation in mouse MII oocytes from three age groups increased with age^73^, further supporting the link between mitochondria function and oocyte quality. Knockout of the BCKD activator *Ppm1k* in female mice leads to increased plasma levels of BCAA and reproductive dysfunction^74^. Additionally, BCAT1 was among the transcripts found to be higher in young human oocytes compared to old (“advanced maternal age”) MII oocytes^75^. Thus, it seems that the mitochondrial regulators of metabolism we have identified here that affect reproductive aging and oocyte quality in *C. elegans* are likely to play conserved roles in the reproductive health of higher organisms, as well.

The genetic pathways that regulate oocyte quality with age are conserved from *C. elegans* to humans^9^, highlighting their fundamental importance in reproductive longevity. Our findings may pave the way for mitochondrial metabolism-based therapeutic interventions, such as Vitamin B1 dietary supplementation, to mitigate the effects of reproductive aging and to promote reproductive health in women.

## Methods

### *C. elegans* genetics and maintenance

Worm strains were provided by the *C. elegans* Genetics Center (CGC). A list of strains used in this study is supplied in Supplemental Table 5. *C. elegans* strains were cultured using standard methods. Worms were maintained at 20°C on plates made from either nematode growth-medium (NGM): 3 g/L NaCl, 2.5 g/L Bacto-peptone, 17 g/L Bacto-agar in distilled water, with 1 mL/L cholesterol (5 mg/mL stock in ethanol), 1 mL/L of 1 M CaCl2, 1 mL/L of 1 M MgSO4, and 25 mL/L of 1 M potassium phosphate buffer (pH 6.0) added to molten agar after autoclaving, or HGM: 20 g/L Bacto-peptone, 30 g/L Bactoagar, and 4 mL/L cholesterol, with all other components same as NGM. Plates were seeded with OP50 *E. coli* for ad libitum feeding. For synchronizing worms, gravid hermaphrodites were exposed to a 15% hypochlorite solution (8.0 mL water, 0.5 mL of 5N KOH, 1.5 mL sodium hypochlorite) to collect eggs, which were washed in M9 buffer (6 g/L Na2HPO4, 3 g/L KH2PO4, 5 g/L NaCl and 1 mL/L of 1M MgSO4 in distilled water) before transferring to seeded plates.

For RNAi experiments, standard NGM molten agar was supplemented with 1 mL/L of 1M IPTG (isopropyl β-d-1-thiogalactopyranoside) and 1 mL/L of 100 mg/mL carbenicillin (RNAi plates). Plates were seeded with HT115 *E. coli* containing the RNAi plasmid or empty control vector (L4440). RNAi clones were obtained from the Ahringer RNAi library and induced with 0.1 M IPTG 1 hour prior to transfer, starting at adulthood. Worms were transferred to freshly seeded plates every other day when producing progeny, unless otherwise specified. For all RNAi experiments, worms were transferred to RNAi plates that were pre-sequence verified.

### Mitochondria purification for mass spectrometry and proteomic analysis

Synchronized *C. elegans* were maintained on HGM agar plates seeded with *E. coli* OP50 as the food source. Every two days, worms were washed off plates with M9 buffer and transferred to fresh plates to avoid progeny accumulation. Mitochondria were isolated using the following optimized protocol modified from previously described methods^76^.

Worms at the appropriate adult stage (as indicated per experiment) were washed off plates with M9 buffer and transferred to HGM agar plates spotted with 5 mg/mL serotonin to induce egg laying. After incubation for 2 hours at 20°C, worms were collected from plates and washed 5x with M9 buffer. Worms were then washed with ice-cold 0.1 M NaCl, pelleted at 1100 x g for 5 minutes at room temperature, and resuspended with an ice-cold mix of 0.1 M NaCl and 60% sucrose (1:1 v/v ratio). Animals were centrifuged at 1100 x g for 5 minutes, and floating worms were collected and diluted 1:4 in 0.1 M NaCl.

Worms were then washed twice with 0.1 M NaCl and resuspended in mitochondria isolation buffer (MIB: 50 mM KCl, 110 mM mannitol, 70 mM sucrose, 0.1 mM EDTA, pH 8.0, 5 mM Tris-HCl, pH 7.4). The resuspended worms were transferred to a cold glass 2 mL Dounce homogenizer (KIMBLE KONTES # 885300-0002) and were homogenized using 20 strokes (B pestle). To remove nuclei and cell debris, the homogenate was centrifuged at 750 x g for 5 min at 4 °C. The supernatant was collected and set aside at 4 °C, while the pellet was resuspended in MIB for an additional round of Dounce homogenization to maximize yields. Following centrifugation at 750 x g for 5 min at 4 °C, the supernatant was pooled with the supernatant collected in the first Dounce step. To remove any residual cell debris and bacteria, the homogenate was filtered through a 5 μm pore size hydrophilic PVDF membrane syringe filter. An aliquot of the homogenate representing cytosolic and membrane cellular fractions was kept for western blot analysis and labeled as the ‘C+M’ sample. The homogenate was centrifuged at 9000 x g for 10 minutes 4 °C. An aliquot of the supernatant representing the cytosolic fraction was kept for western blot analysis, labeled as the ‘C’ sample, and the remaining supernatant was discarded. The resulting pellet (crude mitochondria) was washed with MIB and centrifuged at 10,000 x g for 10 minutes 4 °C. The pellet was resuspended with ice-cold mitochondria resuspension buffer (MRB: 225 mM mannitol, 5 mM HEPES, pH 7.4, 0.5 mM EGTA), and an aliquot of the crude mitochondria was kept for western blot analysis, labeled as the ‘M’ sample. The crude mitochondria fraction was further purified over a Percoll gradient^77,78^; the crude mitochondria suspension was layered on top of a Percoll medium (225 mM mannitol, 25 mM HEPES, pH 7.4, 1 mM EGTA, 30% Percoll (v/v)). Samples were centrifuged at 95,000 x g for 30 minutes using an Optima XE-100 Beckman centrifuge (SW55Ti rotor). 1 mL fractions were collected, and a sample of each was kept for western blot analysis. The pure mitochondria (Mp) were further diluted in MRB and centrifuged at 6,300 x g for 10 minutes to remove residual Percoll. The pure mitochondria pellet was washed and resuspended with MRB. All samples were frozen in liquid nitrogen and kept at -80°C until use.

### Western blot analysis

For western blot analysis, samples were mixed with 10X Bolt Sample Reducing Agent and 4X Bolt LDS Sample Buffer. Samples were heated at 70°C for 10 minutes before loading on a gradient-PAGE (4% -12%) Bis-Tris gel. After their separation, samples were transferred to a PVDF membrane and blocked with 5% milk in TBST (10X TBST: 200 mM Tris-HCl, pH 7.5, 1.5 M NaCl, 1% Tween20). Membranes were incubated with one of the following primary antibodies: anti-Histone H3 (1:5,000 dilution), anti-ATP5A (1 μg/mL), or monoclonal anti-αTUBULIN (0.5 μg /mL). After washing with TBST, membranes were incubated with the corresponding secondary antibody (either goat anti-mouse IgG H&L (HRP), 1:10,000 dilution, or goat anti-rabbit IgG H&L (HRP), 1:10,000 dilution). Membranes were then washed with TBST and imaged using CL-XPosure Film.

### Mass spectrometry and proteomic analysis

Samples were dissolved in 5% SDS (Sigma #05030) and 50 mM TEAB buffer (Sigma # T7408). Trypsin digestion was performed using S-Trap micro spin columns according to the manufacturer’s protocol (*S-Trap*^*TM*^, PROTIFI).

Trypsin-digested samples were dried completely in a SpeedVac and resuspended with 20 μl of 0.1% formic acid, pH 3.0. Two microliters (about 360 ng) were injected per run using an Easy-nLC 1200 UPLC system. Samples were loaded directly onto a 50 cm long 75 μm inner diameter nano capillary column packed with 1.9 μm C18-AQ resin (Dr. Maisch, Germany) mated to metal emitter in-line with an Orbitrap Fusion Lumos (Thermo Scientific, USA). Easy-Spray NSI source was used. Column temperature was set at 45°C, and the two-hour gradient method with 300 nL per minute flow was used (with Buffer A: 0.1% formic acid in water, Buffer B: 0.1% formic acid in 80% acetonitrile and 20% water). The mass spectrometer was operated in data-dependent mode with a resolution of 120,000 for MS1 scan (positive mode, profile data type, AGC gain of 4e5, maximum injection time of 54 sec, and mass range of 375-1500 m/z) in Orbitrap followed by HCD fragmentation in ion trap with 35% collision energy. Charge states of 2 to 7 were included for MS/MS fragmentation. Dynamic exclusion list was invoked to exclude previously sequenced peptides for 60s and maximum cycle time of 3s was used. Peptides were isolated for fragmentation using a quadrupole (1.2 m/z isolation window). Ion-trap was operated in Rapid mode. Raw data were analyzed using the MaxQuant software suite 1.6.6.0^79^ and the Andromeda search engine. The data were queried with the Andromeda search engine against the *C. elegans* and *E. coli* proteome databases appended with common lab protein contaminants. The higher-energy collisional dissociation (HCD) MS/MS spectra were searched against an in silico tryptic digest of *C. elegans* proteins from the UniProt/Swiss-Prot sequence database (v. 2019_09) containing 27,028 sequences, including common contaminant proteins. All MS/MS spectra were searched with the following MaxQuant parameters: acetyl (protein N-terminus), M oxidation; cysteine carbamidomethylation was set as fixed modification; max 2 missed cleavages and precursors were initially matched to 4.5 ppm tolerance and 20 ppm for fragment spectra.

Peptide spectrum matches and proteins were automatically filtered to a 1% FDR based on Andromeda score, peptide length, and individual peptide mass errors. Proteins were identified and quantified based on at least two unique peptides and the label-free quantification (LFQ)^80^ values reported by MaxQuant. The resulting protein groups were imported into Perseus^81^. Data were filtered to include proteins identified with 2 peptides or more and those replicated in at least two of three replicates in at least one group. The data were log2 transformed, and a student’s t-test was used to identify statistically significant proteins. Significant proteins were considered as proteins with more than 1 peptide per protein, fold change >2, and FDR> 0.05. Enrichment analyses were performed using g:Profiler.

### Oocyte fertilizability analysis

Synchronized hermaphrodites were mated with young (day 1) *fog-2* males at L4 stage. After 24 hours, hermaphrodites were singled out and were allowed to reproduce for one day. The number of fertilized embryos (either hatched or unhatched) and unfertilized oocytes produced by each individual were counted daily. *p*-values were calculated using Chi-square test.

### BCAA assay

Assays were performed using the Branched Chain Amino Acid Assay Kit (Abcam, ab83374) according to the manufacturer’s manual with the following modifications. At least 70 worms per condition were Dounce homogenized in 500 μl assay buffer. Reads were normalized to total protein concentration as measured using the Quant-iT Protein Assay Kit. OD was measured at 450 nm using the Synergy MX microplate reader.

### mtROS assay

mtROS assays were conducted according to published protocols. To validate appropriate staining, synchronized adult N2 worms at Day 2 of adulthood were used. Two experimental groups were established: one group was washed and transferred to tubes containing S-basal medium with the addition of MitoTracker Red CM-H2XRos with or without 200 mM of paraquat to induce oxidative stress. The control group was washed and transferred to a tube containing S-basal medium only, without any dye or oxidative stress inducer. All groups of worms were then incubated for 3 hours at room temperature. Following incubation, all worms were gently transferred to NGM plates for a 1-hour recovery phase. Subsequently, the germlines of the worms were dissected and subjected to imaging (Extended Data Fig. 3). For RNAi experiments, synchronized hermaphrodites were transferred to RNAi plates at L4 stage and were transferred to fresh RNAi plates every two days to avoid progeny accumulation. For all other experiments, NGM plates were used. For assays testing Vitamin B1, OP50-seeded NGM plates were used either with or without 5 mg/mL Vitamin B1. One day prior to imaging, worms were transferred to seeded plates with 20 μM MitoTracker Red CM-H2Xros. On the day of imaging (either day 1 or day 5 of adulthood), worms were recovered for 1 hour on seeded RNAi plates without MitoTracker Red CM-H2Xros. Worms were then put on slides with levamisole and dissected for germline imaging. Images were captured on a Nikon eclipse Ti at 60x magnification, focusing on the most mature oocyte (−1 oocyte). All figures shown were from the same experiment, imaged under the same conditions. For each image, an ROI of defined dimensions was drawn around the -1 oocyte. The ROIs were then denoised and deconvolved (using the NIS Elements Denoise.ai and 2D deconvolution functions respectively) for better visualization of subcellular structures. All images in each figure were processed in the same way across the entire image (e.g., adjusting for brightness and contrast).

### TMRE assay

TMRE assays were conducted according to published protocols^82–84^. To validate appropriate staining, synchronized adult N2 worms at Day 2 of adulthood were divided into two groups: one was transferred to culture plates containing Tetramethylrhodamine Ethyl Ester (TMRE) dye to assess mitochondrial membrane potential (ΔΨm), while the control group was placed on plates without dye. On the imaging day, worms from both groups were picked into tubes with S-basal medium, with one subset exposed to 10 μM of FCCP (carbonyl cyanide 4 (trifluoromethoxy)phenylhydrazone) for 1 hour at room temperature to induce mitochondrial membrane depolarization. Afterward, germlines were dissected and imaged, enabling the assessment of the impact of FCCP-induced changes in ΔΨm on oocytes (Extended Data Fig. 3). For TMRE assays, synchronized hermaphrodites were transferred to RNAi plates at L4 stage and were transferred to fresh RNAi plates every two days to avoid progeny accumulation. A 0.5 M stock solution of Tetramethylrhodamine, Ethyl Ester, Perchlorate (TMRE, Invitrogen #T669) was prepared in DMSO and kept at - 20°C. One day prior to imaging, worms were transferred to seeded RNAi plates with 1 mM TMRE (diluted in M9 buffer). On the day of imaging, worms were recovered for 1 hour on seeded RNAi plates without TMRE. For assays testing Vitamin B1, OP50-seeded NGM plates were used either with or without 5 mg/mL Vitamin B1. Worms were put on slides with levamisole and dissected for germline imaging. Images were captured on a Nikon eclipse Ti at 60x magnification, focusing on the most mature oocyte (−1 oocyte). All figures shown were from the same experiment, imaged under the same conditions. For each image, an ROI of defined dimensions was drawn around the -1 oocyte. The ROIs were then denoised and deconvolved (using the NIS Elements Denoise.ai and 2D deconvolution functions respectively) for better visualization of subcellular structures. All images in each figure were processed in the same way across the entire image (e.g., adjusting for brightness and contrast).

### Late-mated reproductive span assay

Late-mated reproductive span assays were performed as previously described^85^. Briefly, hypochlorite-synchronized eggs were placed on plates seeded with OP50 *E. coli*. Starting on L4 stage, ∼25 hermaphrodites were transferred to plates freshly seeded with HT115 *E. coli* containing either empty control vector or RNAi plasmid. Worms were transferred to fresh RNAi plates every two days to avoid progeny accumulation. On the day of late mating, single hermaphrodites and three young *fog-2(q71)* males were transferred to plates seeded with the corresponding HT115 RNAi or control *E. coli* strain. Progeny production was assessed after 4-5 days. All late mating assays were performed at 20°C.

### Full reproductive span

Reproductive span assays were performed either on plates as previously described^13^, or using the lab-on-chip device *Ce*Lab^86^. For both reproductive span assay methods, hypochlorite-synchronized L4 hermaphrodites were mated for 24 h with young adult (day 1) *fog-2(q71)* males at a 1:3 ratio. For assays on plates, each hermaphrodite was moved daily to individual fresh plates until progeny production ceased for two consecutive days. The day of reproductive cessation was counted as the last day of viable progeny production preceding two consecutive days with no progeny. This was assessed by evaluating plates for progeny 2 days after the hermaphrodites had been moved to fresh plates. Successful mating was verified by evaluating the fraction of male progeny produced.

For *Ce*Lab reproductive span assays, chips were fabricated using standard soft lithography to create 200 parallel incubation arenas within each chip. Day 1 worms were loaded into the *Ce*Lab chip as previously described^86^. Animals were incubated in S-medium with OD600 OP50-1, 50 mg/L streptomycin, and 10 mg/L nystatin. For RNAi experiments, bacteria were induced with 4 mM IPTG, pelleted, and resuspended with 2 ml S-medium with 50 mg/L streptomycin, 10 mg/L nystatin, 100 μg/mL carbenicillin and 4 mM IPTG before incubation. Worm survival was scored daily using *Ce*Aid^87^. After each scoring session, progeny were flushed out of the chip using M9 buffer solution for 20 minutes, and the bacteria incubation solution was replenished. Animals were censored, if needed, on the day of loss. Matricide animals were censored the day after. All reproductive spans were performed at 20°C.

For experiments including supplements: all supplements were added to worms starting at day 1 of adulthood. Supplements were added at the appropriate concentration to 10x concentrated heat-killed OP50 *E. coli*. Bacteria were heat-killed by pelleting and resuspending at 10X in double-distilled water. Bacteria were t heated at 65°C for 30 minutes before supplement addition.

### Oocyte quality assay

Oocyte quality assays were performed as previously described^9,18^. For RNAi experiments, synchronized hermaphrodites were transferred to RNAi plates at L4 stage. Worms were transferred to fresh corresponding RNAi plates every two days to avoid progeny accumulation. Hermaphrodites were mated on day 3 of adulthood for 24 h with young adult (day 1) *fog-2(q71)* males at a 1:3 ratio. For all other experiments, OP50-seeded NGM plates were used. For assays testing Vitamin B1, OP50-seeded NGM plates were used with 5 mg/mL Vitamin B1, starting at day 1 of adulthood. Hermaphrodites were mated at L4 stage for 24 h with young adult (day 1) *fog-2(q71)* males at a 1:3 ratio. Following mating, hermaphrodites were transferred to new plates. Mating was confirmed by the presence of male progeny. On the day of imaging, worms were mounted on slides with 2% agarose pads containing levamisole. Images were captured on a Nikon eclipse Ti at 60x magnification. Scoring was performed blindly, and each image was given a score based on the criteria detailed in the relevant figure.

### Lifespan assay

Lifespan assays were performed either on plates as previously described^14^, or using the lab-on-chip device *Ce*Lab^86^. For plate assays, synchronized hermaphrodites were placed on plates at day 1 of adulthood. Hermaphrodites were transferred to freshly seeded plates every other day when producing progeny, and 2-3 days thereafter.

For *Ce*Lab assays, Day 1 worms were loaded into *Ce*Lab chips. Animals were incubated in S-medium with 60 OD600 OP50-1, 50 mg/L streptomycin, and 10 mg/L nystatin.

For RNAi experiments, bacteria were induced with 4 mM IPTG, pelleted, and resuspended with bacteria with 2 ml S-medium with 50 mg/L streptomycin, 10 mg/L nystatin, 100 μg/mL carbenicillin and 4 mM IPTG before incubation. Survival was scored daily using *Ce*Aid^87^. After each scoring session, progeny were flushed out of the chip using M9 buffer solution for 20 minutes, and then the incubation solution was replenished. Worms were censored, if necessary, on the day of matricide, the appearance of abnormal vulva structures, or loss. Animals were defined as dead when they no longer responded to touch. All lifespans were performed at 20°C.

For experiments including supplements: all supplements were added to worms starting at day 1 of adulthood. Supplements were added at the appropriate concentration to 10x concentrated heat-killed OP50 *E. coli* as previously described.

### Germline mitochondria imaging

Worms were synchronized and transferred to RNAi plates at the L4 stage. Hermaphrodites were then maintained on RNAi plates and transferred to fresh RNAi plates every two days to avoid progeny accumulation.

On the day of germline imaging, worms were mounted on slides with 2% agarose pads and levamisole. Images were captured on either a Nikon eclipse Ti at 60x magnification or on a Nikon Ti-2 CSU-W1 spinning disk confocal with SoRa super-resolution technology. For Nikon Ti-2 CSU-W1 spinning disk confocal, a 100X/1.4 NA silicone oil objective was used for imaging, and a Hamamatsu BT Fusion sCMOS (6.5 μm pixel with 93% QE) camera was used for detection. Z stacks were acquired using 561 nm excitation wavelength to image mKate2-labeled mitochondria, and a single slice DIC image of the worm was acquired in the center of the Z stack for reference. Z stacks were processed using the Nikon NIS elements software. 3D volumetric views and movies were created using NIS Elements ‘Volume projection’ and movie maker, respectively.

Scoring was performed blindly. Each image was scored as normal (youthful oocyte characteristics: cuboidal shape, uniform size, organized in the gonad) or abnormal score while focusing on the mature oocytes. Images were processed in FIJI and Adobe Photoshop across the entire Z-stack. For each figure, images shown were from the same experiment, imaged on the same microscope at the same magnification, and with the same exposure. All images in each figure were processed in the same way across the entire image (e.g., max. intensity for Z-stack, adjusting for brightness and contrast).

### Oocyte collection for RNAseq

BA17 or CQ733 worms were synchronized by bleaching and sterilized by incubation at 25°C from L2 to L4. Starting at L4, worms were washed with M9 and transferred to freshly seeded NGM plates every other day. PX627 or CQ730 worms were synchronized and sterilized by bleaching on seeded NGM plates supplemented with 2.5 mL 400 mM indole-3-acetic acid (IAA): Alfa Aesar (#A10556) freshly prepared in ethanol. Worms were maintained on auxin plates until L4 stage. At L4 worms were washed with M9 buffer and were transferred to 0.1 M IPTG-induced RNAi plates, seeded with either L4440- or *bcat-1* RNAi-expressing bacteria. Worms were then washed with M9 and transferred to freshly seeded RNAi plates every other day. On day 8 of adulthood, oocytes were extracted as was previously described^18^, by cutting worms in egg salts buffer, followed by repeated nylon net filtration/centrifugation. Final oocyte pellet was resuspended with 1 mL of Trizol LS and kept at -80°C.

### RNA isolation and RNA sequencing

RNA was isolated from purified oocytes by Trizol-chloroform extraction and cleanup using a RNeasy Micro Kit (Qiagen). Purified total RNA samples were first examined on Bioanalyzer 2100 using RNA 6000 Pico chips to evaluate the integrity and concentration (Agilent Technologies, CA). The poly-A containing RNA transcripts in these samples were converted to cDNA using barcoded oligo-dT primers in the reverse transcription reaction to index each sample following the drop-seq method^88^. The pooled barcoded cDNA samples were purified and amplified by PCR, and then turned into sequencing libraries using the Nextera kit (Illumina, CA) to include only the poly-A tail adjacent 3’ ends of RNA transcripts. These libraries were sequenced on Illumina NovaSeq 6000 S Prime flowcell using the 100 cycle v1.5 kit. Raw sequencing reads were filtered by Illumina NovaSeq Control Software and only the Pass-Filter (PF) reads were used for further analysis. Sequences were mapped to ce11 using RNA STAR, and counts were determined using HTSeq-count and the Ensembl-provided gff file. DESeq2 was then used to determine differential expression.

### Gene Set Enrichment Analysis (GSEA)

DESeq2 was used to identify differentially expressed genes from *daf-2;control(RNAi)* compared to *daf-2;bcat-1(RNAi)* with a FDR of 5% and log2FC > 1. GSEA software (v.4.3.2) was then used to test whether the *daf-2* upregulated genes (*bcat-1*-dependent *daf-2* genes) were significantly enriched in microarray data obtained from Day 1 versus Day 5 oocytes in a *fem-1* background. For GSEA of oocytes from *bcat-1* overexpressing worms compared to *daf-2* oocytes, *daf-2* up and downregulated gene sets were obtained from Templeman et al., 2018^18^. Each gene set was then tested for enrichment among a ranked list obtained from DESeq2 comparing Day 8 oocytes from *bcat-*1-overexpressing (OE) worms compared to control oocytes.

### Microarray analysis

Data from scanned microarrays were loaded onto the Princeton University MicroArray database (PUMAdb) and analyzed as previously described^89^. Genes were filtered for presence in at least 60% of arrays (uncentered correlation, average linking). One-class SAM was used to identify differentially expressed genes and to generate a ranked list of genes expressed Day 1 versus Day 5 oocytes using per gene SAM scores.

### Gene Ontology analysis

Upregulated and downregulated differentially expressed genes from RNA-sequencing experiments were separately analyzed using g:Profiler (version *e109_eg56_p17_1d3191d*).

### Data collection

No statistical methods were used to pre-determine sample sizes but our sample sizes are similar to those reported in previous publications. Data distribution was assumed to be normal but this was not formally tested. Animals were randomized during assignment to experimental groups. Data points were not excluded from any analysis unless noted and explained in the Methods. Except as noted in specific Methods, data collection and analysis were not performed blind to the conditions of the experiments.

### Quantification and statistical analysis

For late-mated reproduction assays, Chi-square tests were used to compare the percentages of worm populations capable of progeny production. Full reproductive span and lifespan assays were assessed using standard Kaplan-Meier log-rank survival tests, with the first day of adulthood of synchronized hermaphrodites defined as t= 0. For oocyte quality and fertilizability experiments, chi-square tests were used to determine if there were significant differences between populations for each category of scored phenotypes. Unpaired t-tests were used for BCAA levels comparisons, for normal/ abnormal germline mitochondria scoring, and for comparing mtROS. For BCAA assays, reads below the detection limit were excluded.

## Data Availability

All RNA-sequencing data are deposited under the PRJNA966212 BioProject. Microarray data are available on the PUMA database at http://puma.princeton.edu as experiment set #7368. Proteomics data are available on PRIDE as Accession #PXD048296. Source data are provided with this paper.

## Code Availability Statement

No custom code was used for this study.

## Acknowledgments

We thank Henry Shwe and Saw Kyin (Princeton Proteomics and Mass Spectrometry Core) for their assistance in sample processing and MS instrumentation, Jennifer Miller and Jean Volmar in the Genomics Core Facility of Princeton University for performing library preparation and next-generation sequencing, Gary Laevsky and Sha Wang (Confocal Imaging Facility, a Nikon Center of Excellence, in the Department of Molecular Biology at Princeton University), The De Botton Protein Profiling institute of the Nancy and Stephen Grand Israel National Center for Personalized Medicine Weizmann Institute of Science (Dr. Yishai Levin), the Petry lab and the Gitai lab (the Department of Molecular Biology at Princeton University) for help with equipment, the *Caenorhabditis* Genetics Center (CGC) for strains, and Murphy Lab members for comments on the manuscript. The study was supported by funding to CTM from the Global Consortium for Reproductive Longevity and Equality (GCRLE) (AWD1006679), The Simons Foundation (811235SPI), The Glenn Foundation for Medical Research, and Pioneer (DP1AG077430) and Transformative (R01AT011963) grants from the NIH.

## Author contributions

Conceptualization, C.L., R.K., and C.T.M.; methodology, C.L., R.K., and C.T.M.; investigation, C.L., J.M.A., R.K., S.S., V.C., T.S., W.K., and S.L.; writing, C.L., R.K., and C.T.M.; funding acquisition, C.T.M.

## Competing Interests Statement

The authors declare no competing interests.

## Extended Data

**Extended Data Figure 1:**
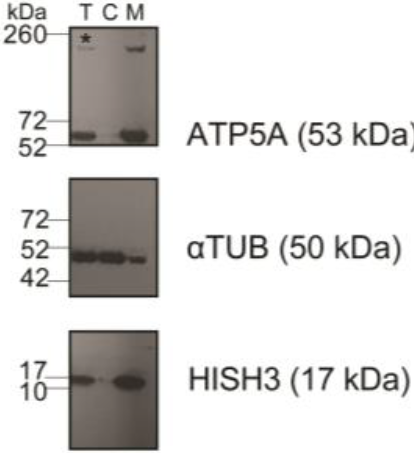
Highly purified mitochondria are isolated from *C. elegans*. Representative western blot of the different cellular fractions. The mitochondrial marker ATP5A is prominently enriched in the membranous fraction, unlike the cytosolic marker aTUB. The membranous fraction was then loaded on a gradient to highly purify the mitochondrial fraction. T-total homogenate, C-cytosolic fraction, M-crude membranes fraction. aTUB-atubulin, HISH3-histone H3. The asterisk represents a non-specific band detected when using the anti-ATP5A antibody. Dashed lines mark where membranes were cut prior to incubation with the indicated antibody. 3 biological replicates were performed.

**Extended Data Figure 2:**
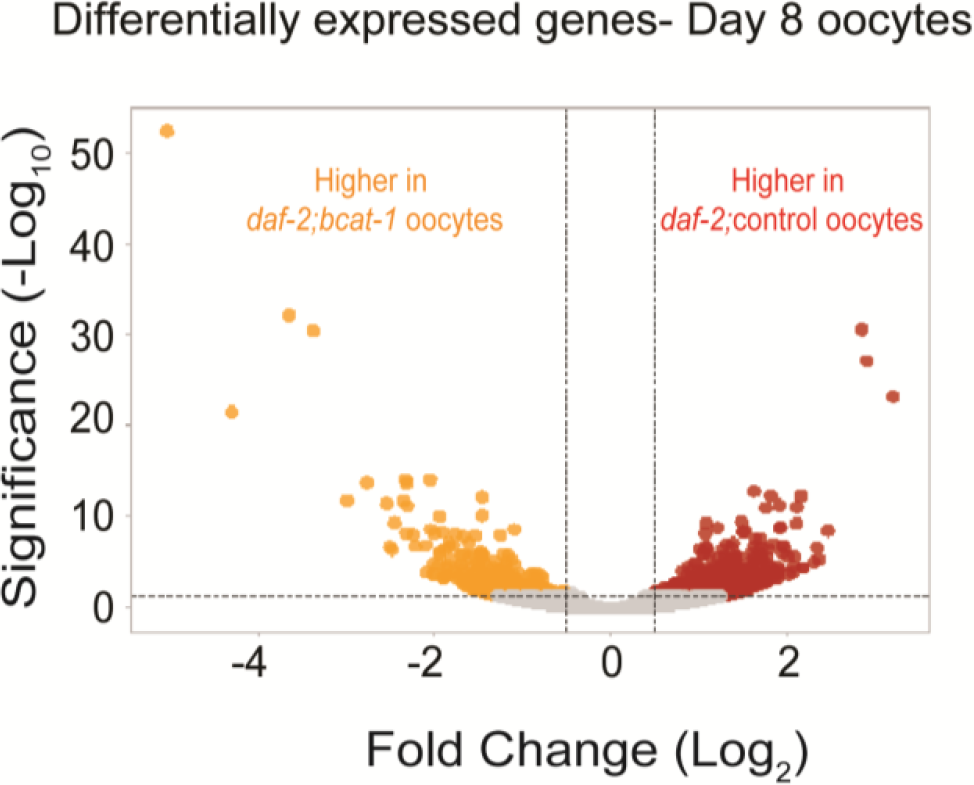
Differentially expressed oocyte genes in day 8 *daf-2*;control and *daf-2;bcat-1(RNAi)* worms. Volcano plot of differentially expressed genes identified between *daf-2*;control and *daf-2;bcat-1(RNAi)* Day 8 oocytes. Red dots denote upregulated genes in *daf-2*;control oocytes, and orange dots denote genes downregulated in *daf-2*;control oocytes relative to *daf-2;bcat-1(RNAi)* (0.5 <log2FC <-0.5).

**Extended Data Figure 3:**
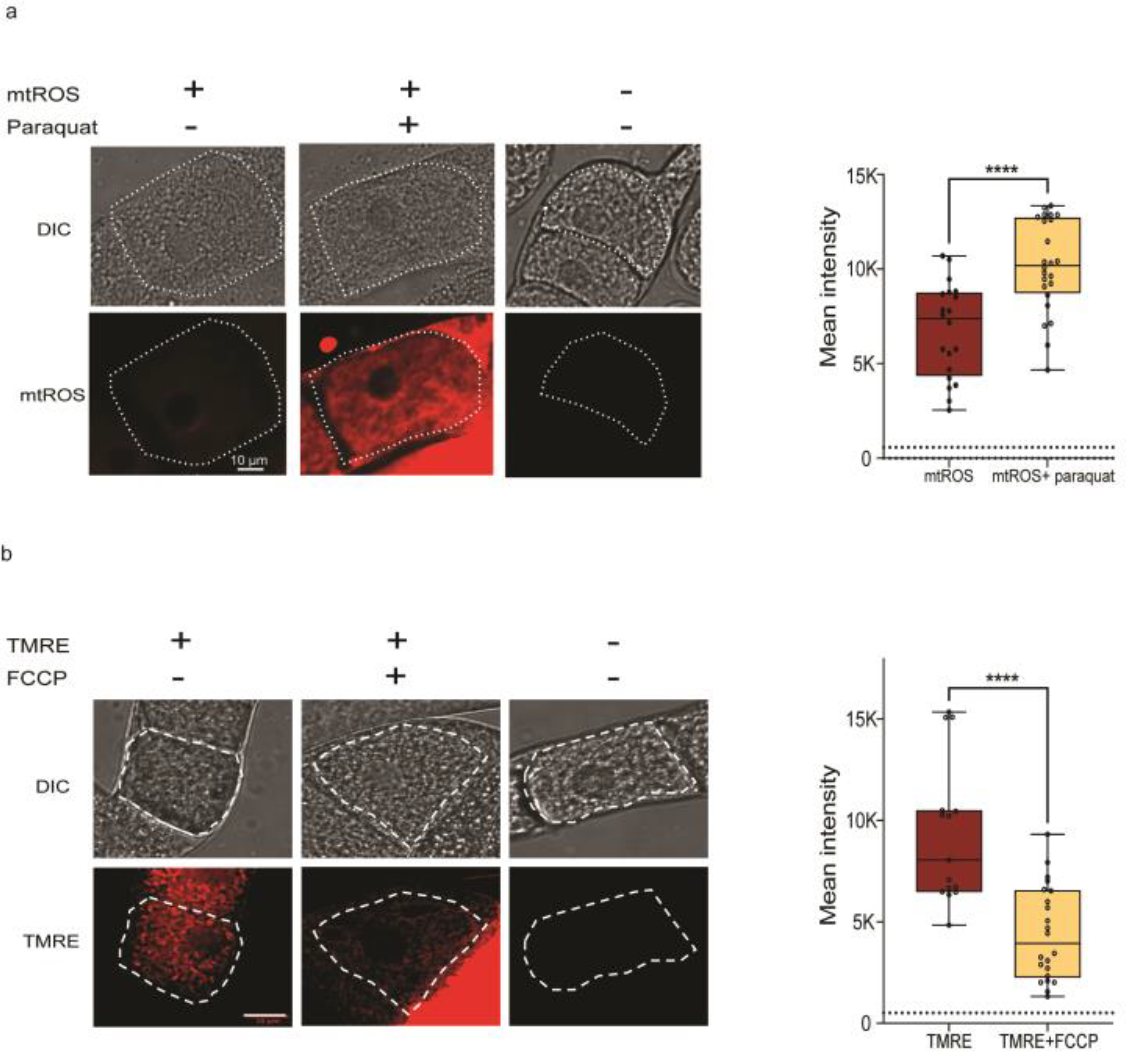
Controls for mitochondrial assays. (a) mtROS levels increase in oocytes after paraquat treatment. A white dashed outline marks the most mature oocytes (−1 oocyte). Right: mtROS levels quantified in mature oocytes either with or without paraquat treatment. Control n = 20, Paraquat n = 24. (b) Left: FCCP treatment reduces mitochondrial membrane potential in oocytes. A white dashed outline marks the most mature oocytes (−1 oocyte). Right: TMRE levels quantified in mature oocytes under control or FCCP treatment conditions. Control n = 15, FCCP n = 22. (a,b) Two-tailed, unpaired t-test. \*\*\*\**p* ≤ 0.0001. Dashed lines on the graphs represent mean intensity of autofluorescence. (a, b) Box plots: center line, median; box range, 25–75th percentiles; whiskers denote minimum– maximum values. 3 biological replicates were performed.

**Extended Data Figure 4:**
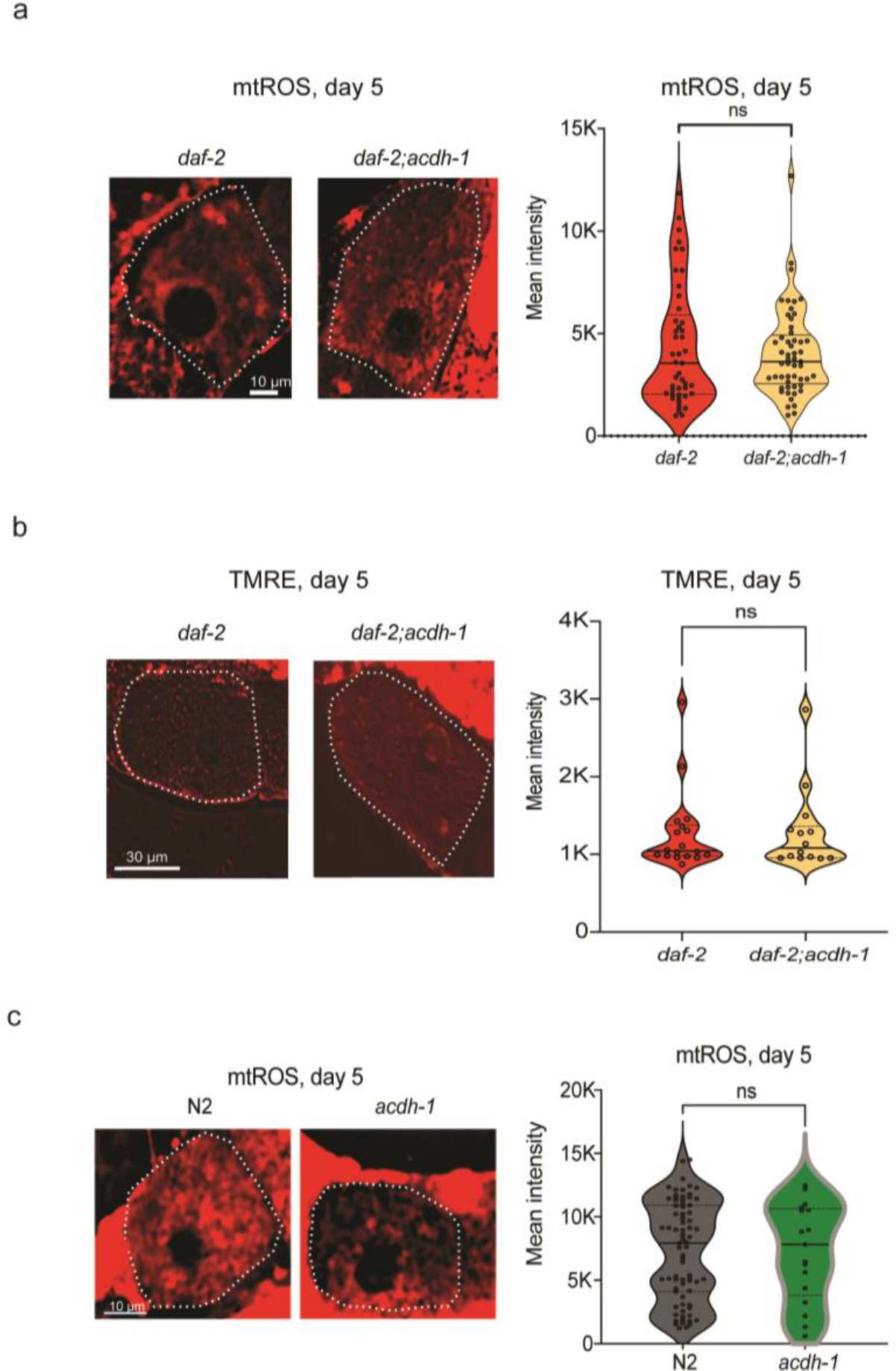
Deletion of *acdh-1* in *daf-2* or wild-type worms does not affect mitochondrial function. (a) Left: mtROS levels do not change in *daf-2* mature oocytes on Day 5 upon *acdh-1* deletion. Right: quantification of mtROS levels in *daf-2;acdh-1* mature oocytes on day 5. N ≥ 45 for each condition. Graph shows results of pooled three biological replicates. (b) Left: *acdh-1* deletion does not affect mitochondrial membrane potential in *daf-2* mature oocytes on day 5. Right: quantification of TMRE signal in *daf-2* mature oocytes on Day 5 upon *acdh-1* deletion. N >10 for all conditions. (c) Left: mtROS levels do not change in wild-type mature oocytes on Day 5 upon *acdh-1* deletion. Right: quantification of mtROS levels in wild-type vs. *acdh-1* mutant mature oocytes on day 5. N ≥ 17 for each condition. Graph shows results of pooled three biological replicates. (a-c) A white dashed outline marks the most mature oocytes (−1 oocytes). Two-tailed, unpaired t-test. ns-nonsignificant. For all panels, representative images are shown. 3 biological replicates were performed.

**Extended Data Figure 5:**
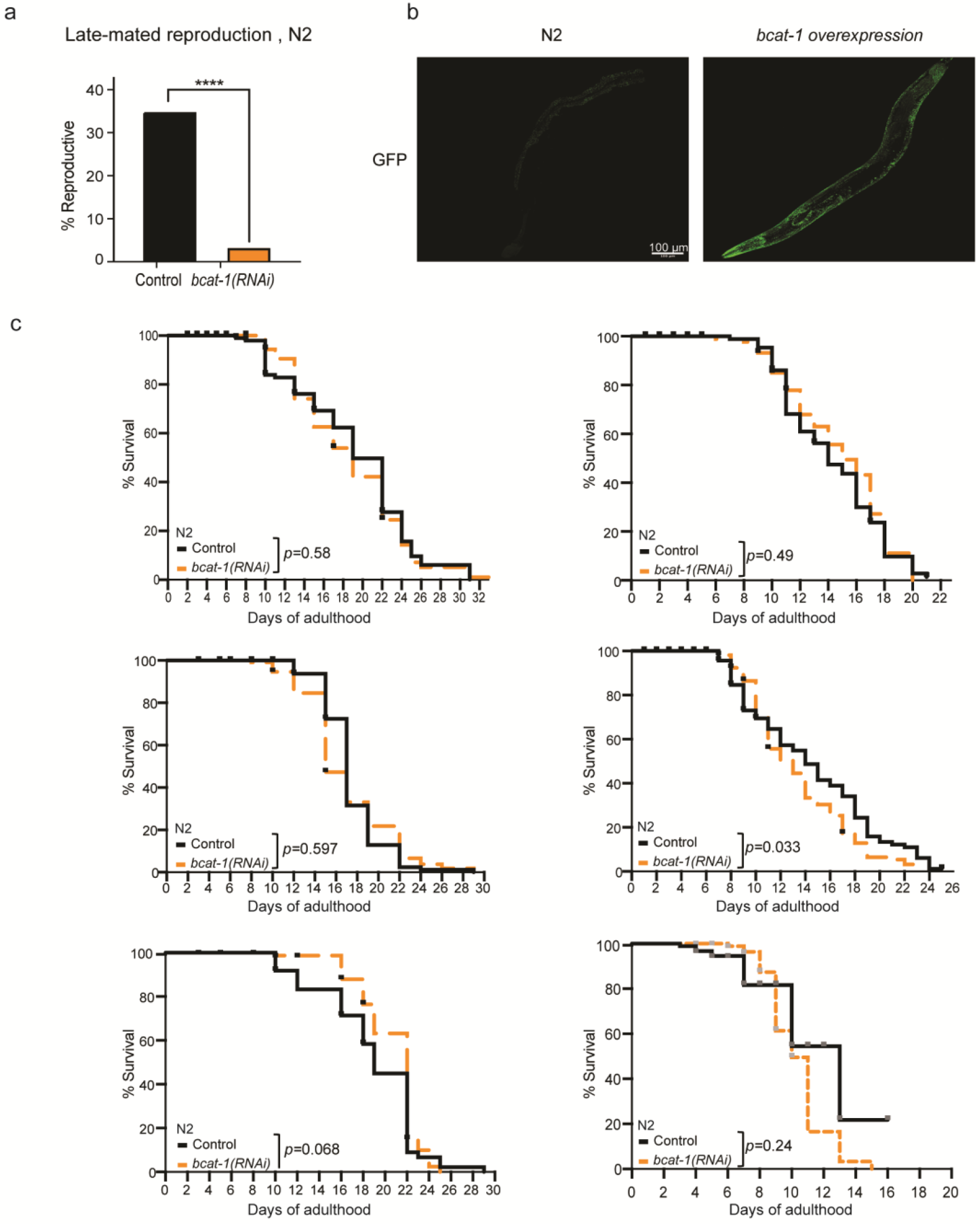
Downregulating *bcat-1* in wild-type worms does not affect lifespan. a) Downregulation of *bcat-1* reduces the late-mated reproduction of adult wild-type worms at day 6 of adulthood. n= 69 for control, n= 58 for *bcat-1(RNAi)*. Chi-square test. \*\*\*\**p* ≤ 0.0001. 3 biological replicates were performed. b) GFP expression in control (left) and in the *bcat-1*-overexpression strain (GFP-BCAT-1 protein fusion). 3 biological replicates were performed. c) Adult-only knockdown of *bcat-1* does not affect the lifespan of wild-type animals. Lifespan data from 6 biological replicates are shown. Results are plotted as Kaplan-Meier survival curves, and the *p-*values were calculated using Mantel-Cox log-ranking. All replicates are shown. Statistical data is detailed in Table S2.

**Extended Data Figure 6:**
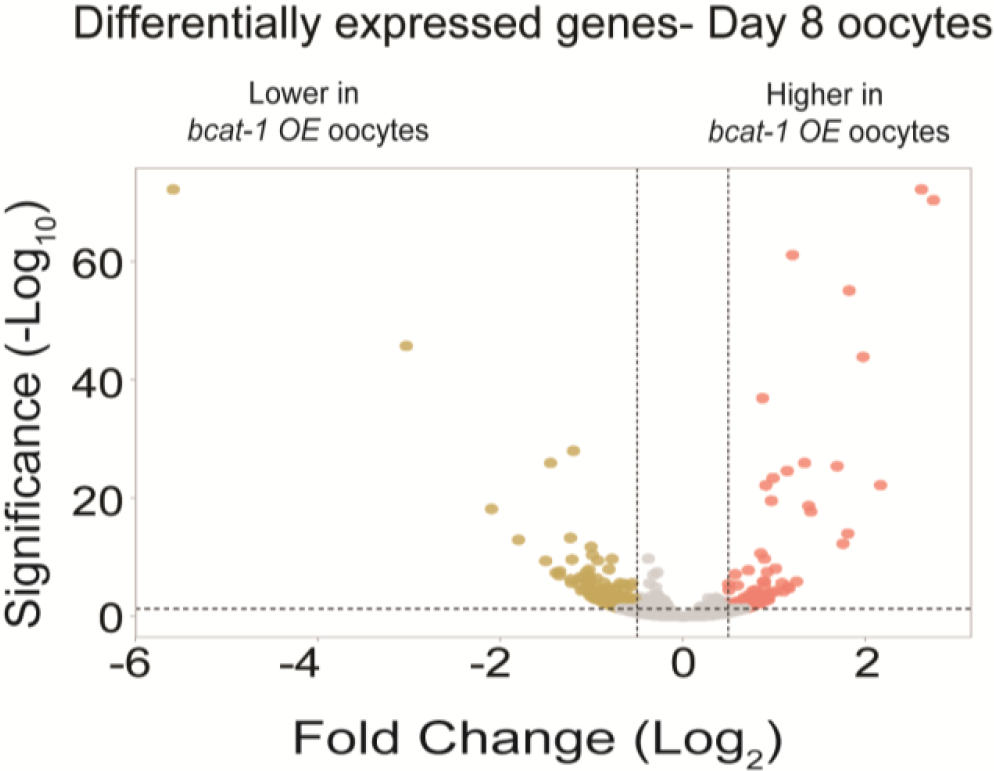
Differentially expressed oocyte genes in day 8 control and *bcat-1* overexpressing worms. Volcano plot of differentially expressed genes identified between Day 8 oocytes from *bcat-1*-overexpressing worms compared to the control (DESeq2, padj < 0.05). Pink dots denote upregulated genes in oocytes from *bcat-1* overexpressing worms, and gold dots denote downregulated genes (0.5 <log2FC <-0.5).

## Notes

### Competing Interest Statement

The authors have declared no competing interest.

### Summary of Updates

Addition of bcat-1 lifespans, Vitamin B1 data

